# Development of DuoMYC: a synthetic cell penetrant miniprotein that efficiently inhibits the oncogenic transcription factor MYC

**DOI:** 10.1101/2024.03.04.583305

**Authors:** Brecht Ellenbroek, Jan Pascal Kahler, Damiano Arella, Cherina Lin, Willem Jespers, Sebastian Pomplun

## Abstract

The master regulator transcription factor MYC is implicated in numerous human cancers, and its targeting is a long-standing challenge in drug development. MYC is a typical ‘undruggable’ target, with no binding pockets on its DNA binding domain and extensive intrinsically disordered regions. Rather than trying to target MYC directly with classical modalities, here we engineer synthetic cell penetrating miniproteins that can bind to MYC’s target DNA, the enhancer box (E-Box), with a high affinity and block MYC-driven transcription. We obtained the miniproteins via structure-based design and a combination of solid phase peptide synthesis and site-specific crosslinking. Our lead variant, DuoMYC, binds to E-Box DNA with high affinity (K_*D*_ = 118 nM) and molecular dynamic simulations provide insights into the structural features leading to the high stability of that DNA binding complex. DuoMYC displays an excellent stability in human serum and is able to enter cells and inhibit MYC-driven transcription with submicromolar potency (IC_*50*_ = 464 nM) as shown by reporter gene assay. Notably, DuoMYC surpasses the efficacy of several other recently developed MYC inhibitors. Our results highlight the potential of engineered synthetic protein therapeutics for addressing challenging intracellular targets.

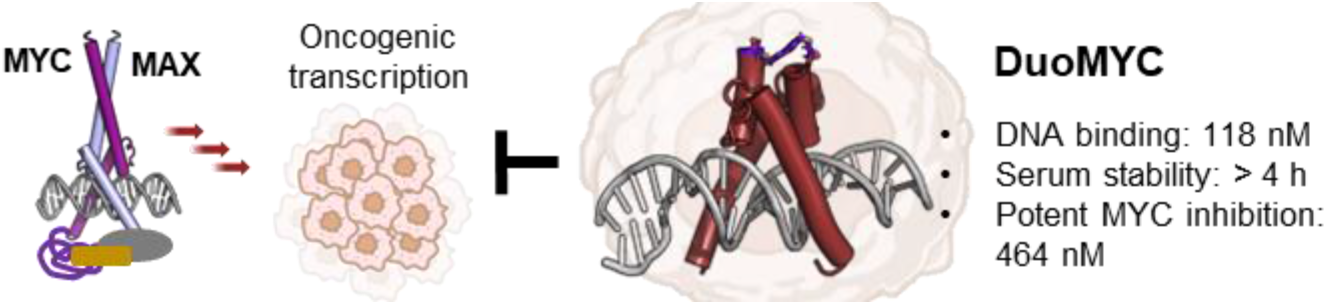

## Introduction

Transcription factors (TFs) orchestrate the delicate equilibrium between gene activation and repression and are crucial for cellular function.^1^ TFs mainly act via protein-protein and protein-nucleic acid interaction and display extensive intrinsically disordered regions used to recruit components of the transcriptional machinery.^2^ These traits make TFs challenging drug targets^3^ and highlight the need for innovative approaches in drug discovery and design.

MYC is a prime example of an undruggable TF. It is involved in the pathomechanisms of over 50% of all human cancers.^4–6^ To exert its activity, MYC heterodimerizes with its partner MAX and as a complex they bind to enhancer box (E-Box) DNA.^7^ MYC’s intrinsically disordered transactivation domain (TAD) then recruits the transcriptional machinery and activates gene programs related to cell growth, proliferation and survival.^8^ MAX can also homodimerize and occupy the same E-Box sequence as a non-productive inhibitory complex.^9–12^ Under healthy conditions MAX/MAX and MYC/MAX are well balanced, while MYC overexpression is involved in many types of cancer. The basic helix-loop-helix leucine zipper motif (bHLH-LZ) of MYC and its intrinsically disordered TAD do not offer any obvious binding sites for inhibitors and no MYC targeted therapeutics are available for patients, to date.

Only recently, a few small molecules with innovative modes of action have shown potential in initial investigations. KI-MS2-008, developed in the Koehler lab, acts as a stabilizer for the MAX/MAX complex, reinforcing this endogenous inhibitor.^13^ EN4, discovered by Nomura and coworkers, is a covalent inhibitor binding to cysteine 171 in the intrinsically disordered domain of MYC and leading to MYC destabilization and degradation.^14^ In addition to these approaches, a MYC-targeting bicyclic peptide was recently discovered from a combinatorial library selection.^15^

An intriguing strategy to target MYC is based on the use of proteins that mimic the inhibitory activity of MAX and occupy E-Box DNA.^16–22^ However, while protein based modalities generally hold immense potential in targeting ‘undruggable’ interactions, their usage is usually limited to extracellular targets, because they are unable to cross cell membranes.^23–25^ Indeed, also the protein-based MYC inhibitors, of which Omomyc is the most studied representant, have mainly been explored as research tools and utilized via ectopic expression in cells or xenografts.^16,18^ However, recently, the Soucek group discovered that purified Omomyc has some intrinsic cell penetrating activity, propelling it from tool compound to a viable therapeutic modality.^26^ An Omomyc variant (Omo-103) is currently being investigated in human clinical trials for the treatment of various forms of tumors.^27^

The intrinsic cell penetration of Omomyc makes it a promising therapeutic modality, but at the same time, cell penetration remains its biggest limitation. Wang *et al*., e.g., found that Omomyc exhibits negligible cellular uptake unless conjugated to a cell-penetrating modality.^28^ These observations correlate well with the fact that, while Omomyc has a low nanomolar binding affinity for E-Box DNA (∼25 nM, see Figure S2), its reported cellular activities are usually 100-1000-fold weaker, when delivered exogenously.^21,22,26,29^

In this study we aim to close this gap and show the development of a synthetic miniprotein that not only has a high affinity for E-Box DNA but, as shown by functional cellular assays, can effectively enter cells and inhibit MYC driven transcription with submicromolar potency.

The MYC/MAX, MAX/MAX, and Omomyc dimers belong to the bHLH-LZ protein family, where the LZ facilitates dimerization, and the basic helix mediates DNA binding. Dimeric configurations are requisite for DNA interaction, whereas monomers lack such affinity. Situated between the basic helix and the LZ, the loop helix domain stabilizes tertiary and quaternary structures, aligning the basic helices appropriately with the major grooves of their target E-Box DNA (Figure 1). Notably, the basic helix domain drives not only DNA binding but also Omomyc’s inherent cell penetration, as evidenced by the loss of this activity upon arginine-to-alanine substitutions within this region.^26^

**Figure 1.**
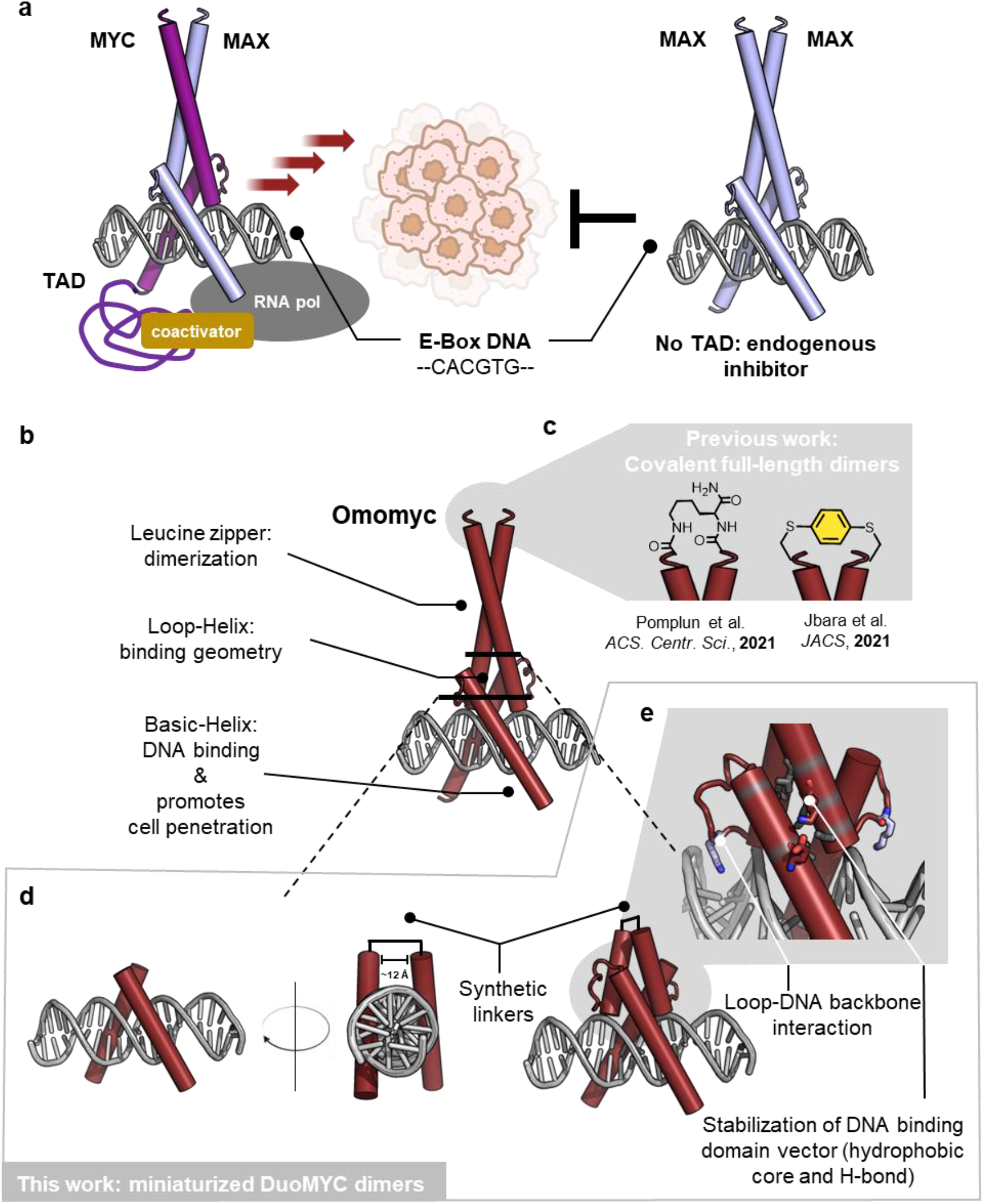
Design of miniaturized protein dimers for MYC inhibition. a) Schematic representation of MYC’s mode of action: MYC upon heterodimerization with MAX binds to E-Box DNA. The MYC-TAD recruits the transcriptional machinery and activates gene transcription. MYC driven gene programs lead to proliferation, cell growth and survival, all common traits of cancer. MAX can also heterodimerize, occupy the same E-Box DNA and act as an endogenous MYC inhibitor. b) Omomyc is a mutated variant of MYC that can also inhibit MYC by occupying E-Box DNA. c) Recent examples of Omomyc variants with covalently linked dimerization domains. d) Design of a miniaturized E-Box binding dimer, consisting of only the two basic helices of MYC. e) Design of a miniaturized E-Box binding dimer, including the loop helix domain for correct and stable orientation of the basic helices into the major grooves of the E-Box sequence.

We reasoned that by miniaturizing Omomyc’s structure, we could create a compact dimeric scaffold with improved cell penetration and biological activity. Given the dual role of the basic helix in cell penetration and DNA binding, we envisioned that the development of synthetically dimerized variants of that portion of the protein could lead to optimized MYC inhibitors. The Moellering group recently showed that miniaturized proteomimetics derived from the MAX bHLH domain can tightly bind to E-Box (3.5 nM) and show improved cell penetration compared to full length MAX.^30^ However, these compounds required > 1000-fold higher concentrations (20 μM) in order to enable significant effects in cellular assays (e.g. reporter gene). Missing a stable dimerization domain, these compounds might not be able to properly dimerize under physiological conditions inside cells. Similarly, compounds dimerized solely via disulfide linkage are likely to be reduced and disassemble inside cells.^29^

Thus, we here hypothesize that in order to obtain compounds with potent cellular activity it is necessary 1) to identify the minimal structure required for specific DNA binding and 2) to covalently link these fragments for a compact and structurally stable scaffold.

## Results and discussion

We analyzed the DNA bound crystal structure of Omomyc^17^ and designed a first miniaturized variant encompassing only the two symmetrical basic helix domains (27 residues). We prepared the dimeric miniMYC (**1**) via parallel automated solid-phase peptide synthesis (SPPS) using lysine (substituted both on the α- and on the ε-amine) as the covalent linkage between the two chains. We first coupled the trifunctional building block Fmoc-Lys(Fmoc)-OH to the solid support and, upon removal of both Fmoc protecting groups, we coupled two glycines and performed a parallel synthesis of the 2 x 27mer sequences (Figure 2a). The Gly-Lys(Gly) linker should effectively bridge the 12 Å distance between the two helices. The dimeric 7 kDa miniMYC (**1**) was obtained in good purity, as shown by HR-UPLC-MS. However, in electro mobility shift assay (EMSA), we detected no binding between E-Box DNA and miniMYC (**1**) (Figure 2b).

**Figure 2.**
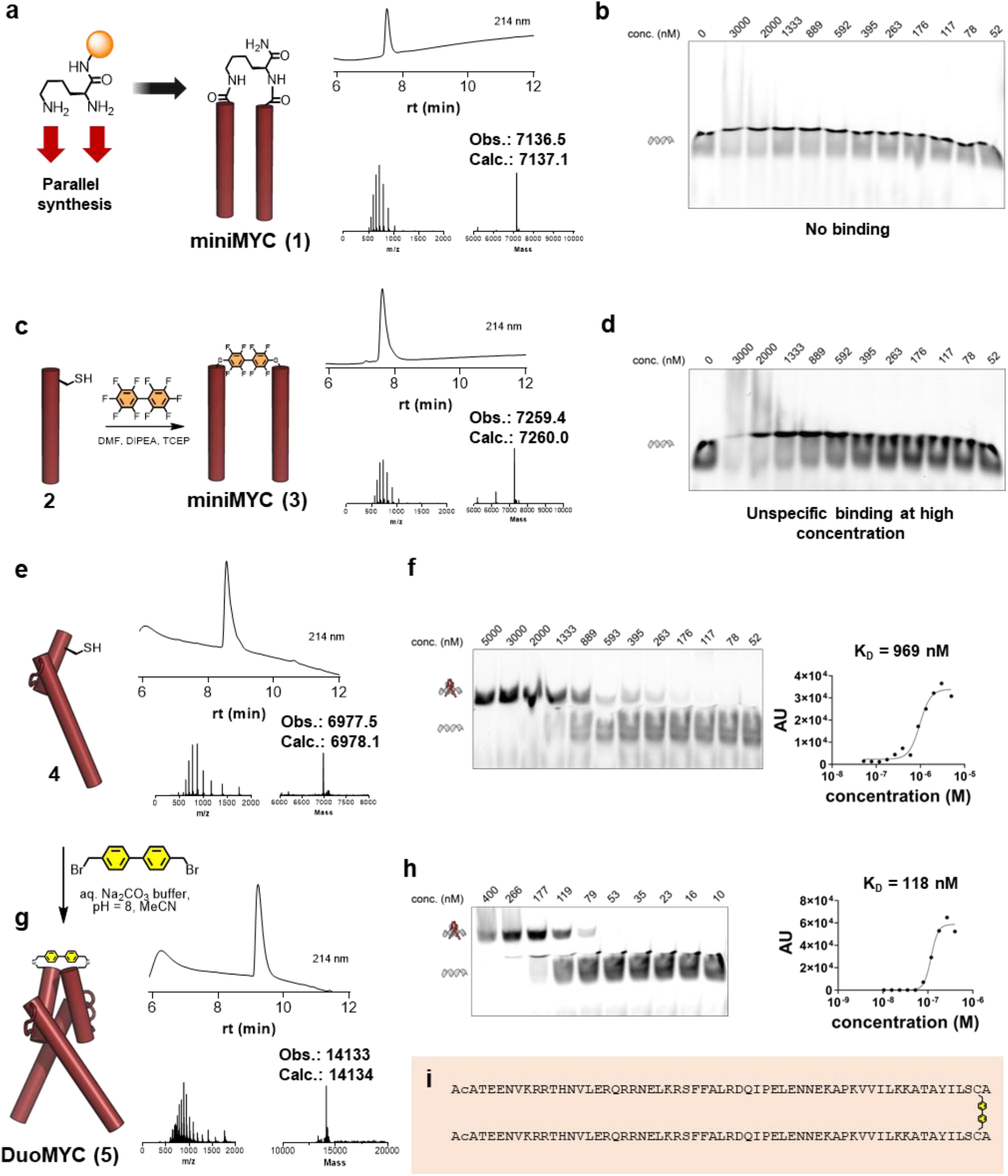
DuoMYC (5) binds to E-Box DNA with nanomolar affinity. a) Parallel synthesis of miniMYC (**1**) via Fmoc-based SPPS with LCMS analysis of the final product. b) EMSA gel of miniMYC (**1**) shows no binding to E-Box DNA. c) Dimerization of miniMYC (**3**) with LCMS analysis. d) EMSA gel of miniMYC (**3**) shows no binding to E-Box DNA. At high concentration unspecific DNA binding was observed. e) SPPS of monomeric variant **4** with LCMS analysis. f) **4** binds to E-Box with a K_D_ = 969 nM. g) Dimerization of **4** with bis(bromomethyl)biphenyl, leading to DuoMYC (**5**). h) DuoMYC (**5**) binds to E-Box with a K_D_ = 118 nM. i) Sequence of DuoMYC (**5**). General SPPS coupling conditions: peptidyl resin incubated with Fmoc-AA-OH (10 eq.), HATU (9 eq.) and DIPEA (29 eq.) in DMF for 8 minutes at 70 °C. Fmoc was removed with 20% piperidine + 2% formic acid in DMF (4 minutes at 70 °C). General EMSA protocol: FAM-labelled dsDNA construct (IRD700-ACCCCAC**CACGTG**GTGCCT, final concentration 4 nM) was preincubated with protein in EMSA buffer (20 mM HEPES, pH 8.0, 150 mM NaCl, 5% glycerol, 1 mM EDTA, 2 mM MgCl_2_, 0.5 mg/mL of BSA, 1 mM DTT and 0.05% NP-40) for 30 minutes at rt followed by 15 minutes on ice and subsequently run on a 10% acrylamide TBE gel under native conditions.

We sought to increase the rigidity of the linker for a variant with a stabilized structure, hoping to improve the binding to E-Box DNA. We synthesized monomeric monomer (**2**) with a Cys close to the N-terminus and linked two monomers using a decafluoro biphenyl reagent. The dimerization reaction worked efficiently and we isolated pure miniMYC (**3**) (Figure 2c). Unfortunately, also this variant did not show any specific association to E-Box DNA (Figure 2d).

We hypothesized that the helix loop domain plays a crucial role in orienting the basic helices into the E-Box major grooves in the correct angle. A parallel synthesis of this medium sized analog failed (data not shown). Therefore, we synthesized a monomeric variant of the 59mer with a single cysteine close to the C-terminus (**4**) (Figure 2e). Encouragingly, this medium-sized variant, without a covalent dimerization linker, exhibited E-Box DNA binding with an affinity of 969 nM (Figure 2f). The loop helix domain might thus drive protein dimerization, at least to some extent. Recognizing, however, that the full-length Omomyc dimerization is mainly driven by the leucine zipper domain which we removed, we again decided to introduce a covalent linker for enhanced structural reinforcement. We cross-linked the two monomers with a bis(bromomethyl)biphenyl linker and successfully generated the synthetic 14 kDa miniprotein dimer, DuoMYC (**5**) (Figure 2g). Notably, DuoMYC (**5**) displayed improved E-Box DNA binding, with a K_*D*_ of 118 nM, representing an 8-fold enhancement over the monomeric variant (Figure 2h). We also prepared a second variant, DuoMYC (**7**), using a decafluorobiphenyl reagent for dimerization, and obtained a K_*D*_ of 174 nM (see Figure S1 for synthesis details and EMSA).

We next compared the stability of the full length Omomyc/Omomyc-DNA complex with the DuoMYC (**5**) DNA complex using molecular dynamic (MD) simulations. The MD simulation was performed using Schrödinger’s Desmond using OPLS4 under the NPγT ensemble. Each system was run in triplicates of 500 ns. We found that both Omomyc and DuoMYC (**5**) remain in a stable DNA bound conformation in each of the three replicate systems, with overall residue fluctuations between 2.2 and 3.5 Å compared with their starting coordinates (Figure 3a). The root-mean-square-fluctuation (RMSF) plot evidences the structural rigidity of the two helical domains in DuoMYC (**5**), while the loop connecting the basic helix to the C-terminal helix displays significantly stronger fluctuations (Figure 3b). A very similar trend could be observed in Omomyc (Figure S9). The flexibility of the C-terminal extremities of the DuoMYC (**5**) dimer enables the correct arrangement of the covalent biphenyl linker, connecting the two helices (Figure 3c-d). Overall, the MD simulation highlights how the compact covalent linker (mw < 200) can efficiently replace the extensive leucine zipper domain (mw > 7000), still resulting in a stable DNA binding scaffold.

**Figure 3.**
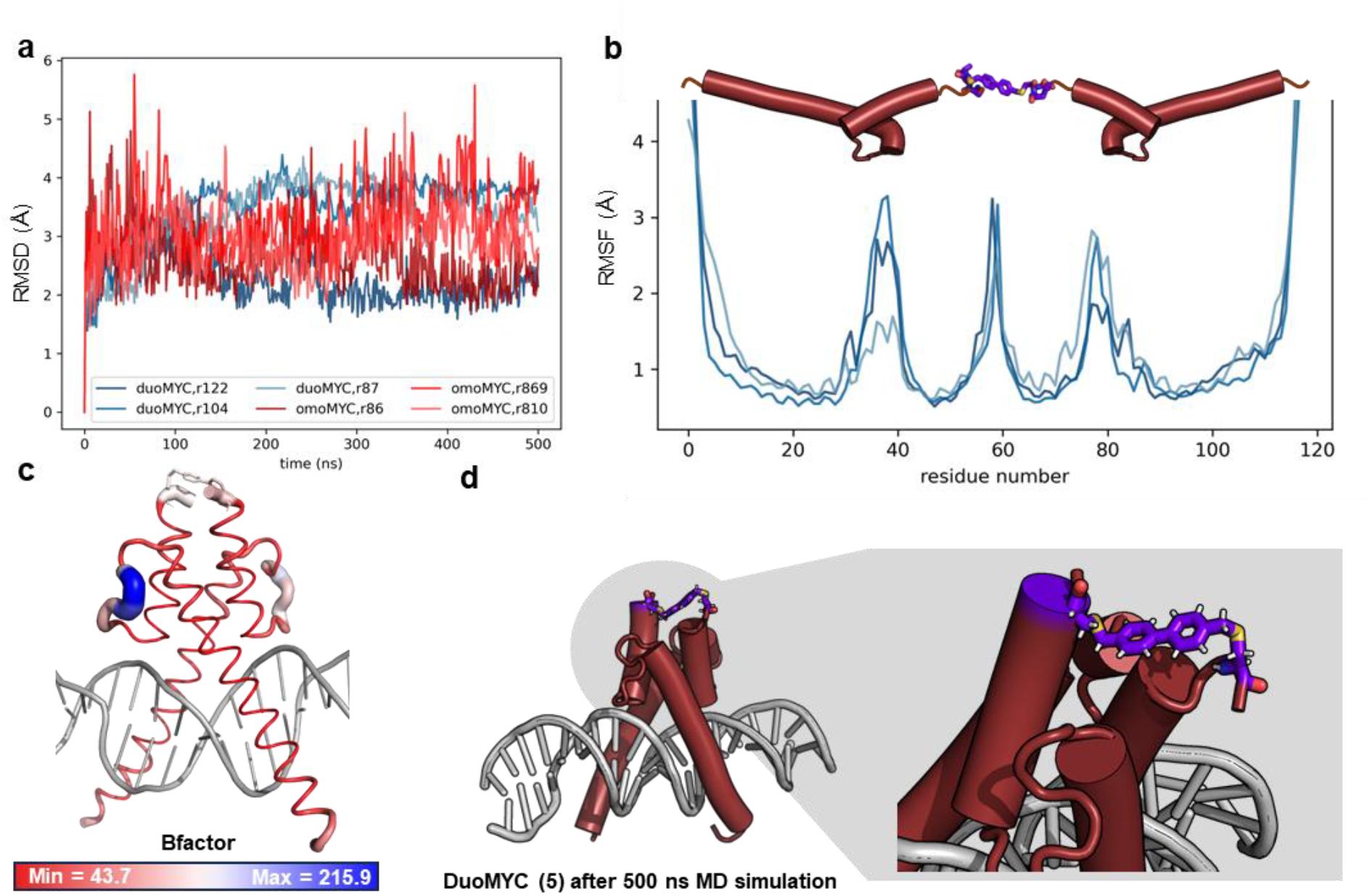
MD simulation shows stability of DuoMYC (5) DNA complex. a) Root-mean-square-deviation (RMSD) plot of DuoMYC (5) and Omomyc, in blue and red, respectively. Each triplicate simulation was run for 500 ns. b) RMSF plot for DuoMYC. c) RMSF plot translated to b-factor and shown as putty representation. d) DuoMYC structure after 500 ns MD simulation with linker region highlighted.

Since proteolytic instability is one of the major limitations of peptide-based therapeutics we next assessed the half-life of monoMYC (**4**) and DuoMYC (**5**) in human serum. We incubated both compounds in human serum and measured their stability by LC-MS at different timepoints (normalizing to an internal standard). MonoMYC (**4**) was rapidly degraded with an overall half-life of 24 minutes. DuoMYC (**5**) in turn, presented an excellent stability with > 50% compound still intact after 4 hours (Figure 4a).

**Figure 4.**
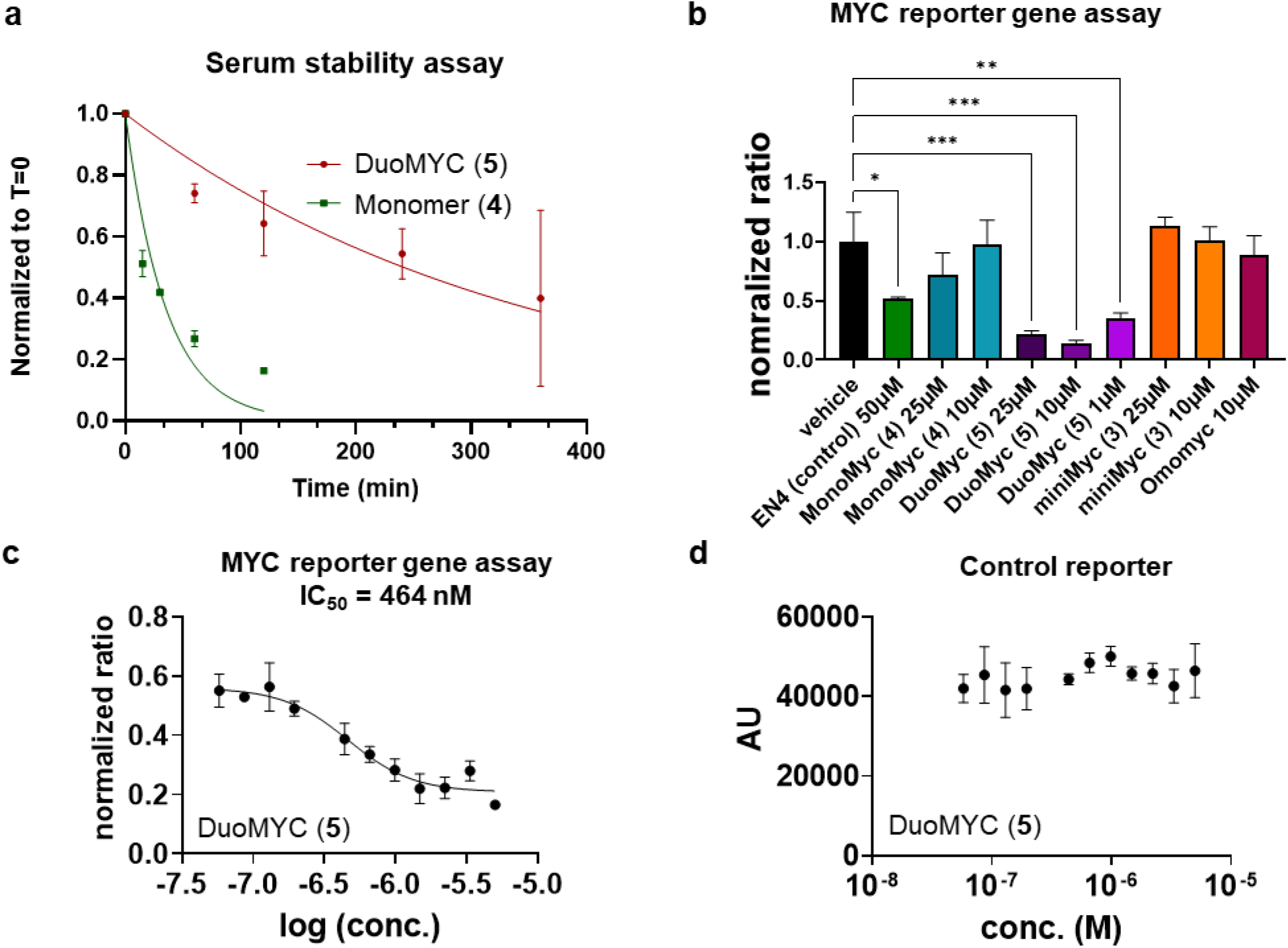
DuoMYC (5) inhibits MYC driven transcription with submicromolar potency and has a good serum stability. a) Monomer (**4**) displays a half-life of 24 min and DuoMYC (**5**) a half-life of 241 min. General serum stability protocol: proteins were dissolved in 10% human serum in DPBS to a final concentration of 60 μM, mixed and after the indicated time aliquots were taken and inactivated with 20% TFA in H_2_O.The samples were analyzed via high-resolution LC-MS and curves plotted in Graphpad Prism 9. The standard deviations plotted are from n = 2 independent samples. b) MYC reporter gene assay with single concentrations of several compounds. HEK293 cells were transfected with a dual reporter gene, containing a MYC dependent firefly luciferase gene and a MYC independent, constitutively expressing renilla luciferase construct. The cells were treated with compound at the indicated concentration for 24 h. Subsequently, the cells were lysed, luciferase luminescence measured and the data were normalized. Statistical significance details: A one-way ANOVA revealed that there was a statistically significant difference in MYC-responsive luminescence between at least two groups F (9, 14) = 12.49, *p* < 0.0001. Dunnett’s multiple comparisons test was performed to test for a significant difference of the various groups from vehicle. * *p*<0.05, ** *p*<0.01, *** *p*<0.001. c) Concentration curve of DuoMYC (**5**) reporter gene activity. d) Control reporter remains stable upon DuoMYC (**5**) treatment.

To assess whether the tight binding activity and the miniaturized, stable structure of DuoMYC (**5**) would translate to improved biological *in cell* activity we performed a MYC reporter gene assay. The reporter is based on a firefly luciferase gene activatable by MYC/MAX-dimers. A second constitutively-expressing Renilla luciferase vector serves as an internal positive control for normalization. HEK293 cells were transfected with the reporter gene DNA and treated with a number of different compounds for 24 h (Figure 3a). As a positive control for MYC inhibition we included the recently discovered covalent small molecule MYC inhibitor EN4, that had shown activity in the same assay format. We also included recombinant Omomyc (see SI 4.2-4.6 for expression protocol and Figure S2 for EMSA validation), miniMYC **3**, monoMYC **4**, and finally DuoMYC. Only EN4 and DuoMYC (**5**) treatments resulted in statistically significant reduction in MYC dependent transcription (Figure 4b). While 50 μM EN4 reduced MYC transcription by ∼ 50%, DuoMYC (**5**) showed stronger inhibition even at 50-fold lower concentration (1 μM). Omomyc did not show any effect on MYC transcription. We next measured a full concentration curve of DuoMYC (**5**) and determined an EC_50_ of 464 nM (Figure 4c). The MYC-independent control reporter showed a stable signal over the whole concentration range of DuoMYC (**5**), indicating that the compound is likely specifically inhibiting MYC driven transcription, rather than non-specifically shutting down cellular gene transcription (Figure 4d).

## Conclusion

Taken together, here we have shown the development of a synthetic cell penetrating miniprotein that can bind E-Box DNA with a high affinity and inhibit MYC-driven oncogenic transcription. Although MYC is a long sought after drug target, often described as undruggable, classical small molecule drugs cannot efficiently target MYC, because of its lack of binding pockets while protein drugs have limited activity due to their poor cell permeability. Our engineered miniprotein DuoMYC (**5**) combines a strong binding affinity and potent activity in cell assays. Furthermore DuoMYC (**5**) exhibits excellent serum stability. We rationally engineered DuoMYC (**5**) based on the crystal structure of a larger E-Box targeting modality, Omomyc.^17^ Surprisingly, our initial efforts to miniaturize the structure toward a dimeric miniprotein containing only the DNA binding helices of Omomyc was unsuccessful while similar approaches had shown success in miniaturized GCN4 mimetics.^31,32^ The mandatory components for our lead compound include the DNA binding helices, the loop helix domains and a stable covalent linkage. This scaffold ultimately resulted in an efficient MYC inhibitor. Given the interest in MYC as a cancer drug target,^33^ DuoMYC (**5**) can be considered a highly promising modality. Furthermore, our results highlight the possibility of designing innovative chemical modalities, such as synthetic miniproteins, that can penetrate cells and address challenging targets.

## ACKNOWLEDGMENTS

BE, JPK and SJP acknowledge funding from the ERC StG (SynTra - 101039354). The Pomplun Lab gratefully acknowledges support from the Oncode Institute and in particular the generous donation from Mr. H.J.M. Roles. We also acknowledge the Leiden Institute of Chemistry protein expression facility for support and the group of Prof. Dr. Eilers at University of Wuerzburg for donating the plasmid used for the recombinant expression of Omomyc.

