## Supplementary information for "Development of DuoMYC: a synthetic cell penetrant miniprotein that efficiently inhibits the oncogenic transcription factor MYC"

**Contents**

List of abbreviations ... 3
1. Reagents and solvents ... 4
2. Synthesis procedures ... 4
2.1 Automated solid phase peptide synthesis (ASPPS) – general protocol ... 4
2.2 Acetylation ... 5
2.3 Full cleavage ... 5
2.4 Preperative HPLC (Prep-HPLC) – General protocol ... 5
2.6 LC-MS ... 5
2.6.1 Method A (low resolution) ... 5
2.6.2 Method B (high resolution) ... 5
2.7 Plasma stability assay ... 6
3 Protein synthesis with LCMS spectra ... 6
3.1 MiniMyc with lysine linker (1) ... 6
3.2 MiniMyc monomer (2) ... 7
3.3 MiniMyc with octafluorobiphenyl linker (3) ... 9
3.4 MonoMyc (4) ... 10
3.5 DuoMyc with 4,4'-dimethyl-1,1'-biphenyl linker (5) ... 11
3.6 MonoMyc (6) ... 13
3.7 DuoMyc with octafluorobiphenyl linker (7) ... 14
4. Biochemistry ... 16
4.1. Overview of buffers used ... 16
4.2. Expression of Omomyc in E.Coli ... 16
4.3. Purification of his-tagged proteins ... 16
4.4. General protocol for SDS-PAGE ... 16
4.5. General protocol for buffer exchange ... 17
4.6. His-tag cleavage by enterokinase ... 17
4.7. Electromobility shift assay (EMSA) ... 17
4.8. Cell culture ... 17
4.9. Reporter gene assay ... 17
5. Molecular Dynamic simulations ... 18
6. Supplementary figures ... 19
Figure S1. ... 19
Figure S2. ... 20
Figure S3. ... 21
Images of uncut gel files ... 22

Plasmide and protein sequences........................................................................................................ 28

Table S1. ....................................................................................................................................... 30

**List of abbreviations**

| | |
|---|---|
| MeCN | Acetonitrile |
| ASPPS | Automated solid-phase peptide synthesis |
| AUC | Area under the curve |
| CV | Column volume |
| DCM | Dichloromethane |
| DIPEA | N,N-Diisopropylethylamine |
| DMF | N,N-Dimethylformamide |
| DPBS | Dulbecco's phosphate buffered saline |
| EDTA | Ethylenediaminetetraacetic acid |
| EK | Enterokinase |
| EMSA | Electrophoretic mobility shift assay |
| HATU | Hexafluorophosphate Azabenzotriazole Tetramethyl Uronium |
| HPLC | High-performance liquid chromatography |
| IS | Internal standard |
| LC-MS | Liquid chromatography-mass spectrometry |
| MD | Molecular dynamics |
| NEAA | Non-essential amino acid |
| PBS | Phosphate buffered saline |
| PDB | Protein data bank |
| RT | Room temperature |
| RP-FC | Reverse phase flash chromatography |
| SDS-PAGE | Sodium dodecyl sulphate polyacrylamide gel elctrophoresis |
| SPPS | Solid-phase peptide synthesis |
| TBE | Tris borate edta buffer |
| TCEP | Tris(2-carboxyethyl)phosphine |
| TFA | Trifluoroacetic acid |
| TIC | Total-ion chromatogram |
| XIC | Extracted-ion chromatogram |

### 1. Reagents and solvents

All chemicals and solvents were directly used as they were purchased. The Fmoc-protected amino acids (Fmoc-Ala-OH, Fmoc-Arg(Pbf)-OH, Fmoc-Asn(Trt)-OH, Fmoc-Asp(OtBu)-OH, Fmoc-Cys(Trt)-OH, Fmoc-Glu(tBu)-OH, Fmoc-Gln(Trt)-OH, Fmoc-Gly-OH, Fmoc-Ile-OH, Fmoc-Leu-OH, Fmoc-Lys(Boc)-OH, Fmoc-Met-OH, Fmoc-Pro-OH, Fmoc-Phe-OH, Fmoc-Ser(tBu)-OH, Fmoc-Thr(tBu)OH, Fmoc-Trp(Boc)-OH, Fmoc-Tyr(tBu)-OH, Fmoc-Val-OH) and 4,4'-Bis(bromomethyl)biphenyl (≥97.0% (HPLC)), Tris(2-carboxyethyl)phosphine hydrochloride (TCEP), acetic acid (≥99.0%), ammonium bicarbonate, Dulbecco's phosphate buffered saline (DPBS) and human serum (from human male AB plasma, USA origin, sterile-filtered) were bought from Sigma Aldrich Merck Milipore. Fmoc-His(Boc)-OH was bought from Carbolution. Fmoc-Lys(Fmoc)-OH was bought from Bachem. For SPPS we used H-rink-amide chemmatrix-HL (0.41 mmol/g). O-(7-azabenzotriazol-1-yl)-N,N,N',N'-tetramethyluronium hexafluorophosphate (HATU, 98%) was purchased from Fluorochem, N,N-Diisopropylethylamine (DIPEA, ≥99 %) and piperidine (≥99 %) were bought from Carl Roth. P-Cresol (99%) and Thioanisole (99%) were purchased from Fisher scientific. Peptide grade N,N-Dimethylformamide (DMF) and acetonitrile (MeCN) (HPLC and LCMS grade) were purchased from Biosolve Chimi. Acetic anhydride (97%) was purchased from Acros organics. Dichloromethane (DCM) was bought from Honeywell. Dietheyl ether was bought from VWR chemicals. Trifluoroacetic acid (Peptide Grade) (TFA) was purchased from Iris Biotech.

$H_2O$ used in the experiments was Mili-Q water obtained from Milli-Q® Reference bought at Avantor and will from this point on be mentioned as $H_2O$.

### 2. Synthesis procedures

#### 2.1 Automated solid phase peptide synthesis (ASPPS) – general protocol

All peptides were prepared on a Syro I XP synthesizer. The synthesis was performed on Chemmatrix rink amide resin (typically 100 mg, loading capacity of 0.41 mmol/g, 41 µmol scale). The synthesizer was charged with Fmoc-protected amino acid (Fmoc-AA-OH) solutions (0.5 M in DMF), HATU (0.45 M in DMF), Fmoc removal cocktail (20%/2%/78% = piperidine/formic acid/DMF) and pure DIPEA. As the first step in the peptide synthesis, the peptidyl resin was incubated at 70 °C in DMF while shaking for 10 min, after which the following coupling cycle was repeated until completion of the synthesis:

**Coupling cycle**

Coupling: Fmoc-AA-OH (10 eq., 800 µL for the 41 µmol scale), HATU (9 eq., 800 µL for the 41 µmol scale) and DIPEA (28.7 eq., 100 uL for the 41 umol scale) were sequentially added to the peptidyl resin and incubated at 70 °C for 8 minutes, with vigorous interval vortexing, followed by vacuum based draining of the resin.

Washing: DMF (1.2 mL) was added to the peptidyl resin, incubated for 1 min at room temperature (RT) with vigorous vortexing followed by vacuum based draining of the resin. The step was repeated 3 times.

Fmoc removal: Fmoc removal cocktail (1.5 mL) was added to the peptidyl resin incubated at 70 °C for 4 minutes, with vigorous interval vortexing.

Washing: DMF (1.2 mL) was added to the peptidyl resin, incubated for 1 min at RT with vigorous vortexing followed by vacuum based draining of the resin. The step was repeated 3 times.

After completion, the peptidyl resin was washed with DMF (3x), DCM (3x), dried under vacuum and either stored at -20 ˚C or directly acetylated.

#### 2.2 Acetylation

After completion of the ASPPS, the peptides were acetylated. First, the dry peptidyl resin was swollen in DMF (10 min) and subsequently vacuum-based drained. To the DMF soaked peptidyl resin the acetylation mixture (10%/10%/80% = $Ac_2O$/DIPEA/DMF) was administered (3 mL for 41 µL scale). The resin was thorougly shaken for 15 min at RT. The resin was subsequently washed with DMF (3x), DCM (3x), dried under vacuum and either stored at -20 ˚C or cleaved immediately.

#### 2.3 Full cleavage

The peptide was cleaved for 2 h using ~10 mL of 'special reagent K' cleavage cocktail (82.5%/5%/5%/5%/2.5% = TFA/$H_2O$/Cresol/Thioanisole/1,2-ethanedithiol). TFA was evaporated under a gentile nitrogen stream. Subsequently, the peptide was triturated (2x) using ice cold diethylether, spun down for 5 min at 10000 RPM. The pellet was dissolved in $H_2O$ (10 mL for 41 µmol scale) and lyophilized.

#### 2.4 Preperative HPLC (Prep-HPLC) – General protocol

For prep-HPLC a BESTA-Technik system attached to a Reprosil Gold 120 C18 column 10 µm (250 x 25 mm) and an ECOM Flash 10 DAD 800 UV detector set on 214 nm was used**.** The mobile phases used were solvent A (94.9%/5%/0.1% = $H_2O$/ MeCN /TFA) and solvent B (5%/94.9%/0.1% = $H_2O$/ MeCN /TFA).

Method: 0% solvent B over 3 min, next 0% to 15% solvent B over 1 min, followed by a linear gradient 15% to 90% solvent B over 86 min and 100% solvent B over 3 min. The obtained fractions where analyzed using a Sciex X500b QTOF LC-MS (see LC-MS high resolution protocol, method B). Pure fractions were combined and stored as lyophilized powders at -20 °C.

### 2.6 LC-MS

LC-MS chromatograms and associated mass spectra were acquired using a Shimazu LCMS-2020 system (Method A) or, for high resolution mass spectrometry data, a Sciex X500b QTOF ESI-QToF mass spectrometer couple to a Shimadzu Nexera UHPLC LC40DX3 (Method B). Mobile phases used for LC-MS analysis are solvent A (0.1% formic acid in $H_2O$) and solvent B (0.1% formic acid in acetonitrile). Solvent C (94.9%/5%/0.1% = $H_2O$/ MeCN /TFA) was used to dissolve protein mixtures to a concentration of 1 mg/mL.

The following LCMS methods were used:

##### 2.6.1 Method A (low resolution)

Column: Kinetex® 2.6µm XB-C18 100 Å LC Column (50 x 3 mm) UV detector: 214 nM.

LC Method: 0% solvent B over 1 min, followed by a linear gradient 0% to 70% solvent B over 10 min, followed by 70% solvent B over 1.5 min, followed by 70% to 0% solvent B over 4.5, flowrate 0.55 mL/min.

##### 2.6.2 Method B (high resolution)

Column: Phenomenex Synergi™ 4 µm Fusion-RP 80 Å LC Column (50 x 2 mm).

LC Method: 0% solvent B over 1 min, followed by a linear gradient 0% to 60% solvent B over 3.5 min, followed by a linear gradient 60% to 95% solvent B over 0.1 min, followed by 95% solvent B over 0.4

min, followed by a linear gradient 95% to 0% solvent B over 0.5 min, followed by 0% solvent B over 1.5 min, flowrate 0.5 mL/min.

MS parameters: General parameters: Method duration: 5 min; Total scan time: 0.276 sec; Estimated cycles: 1086; Intact protein mode: False; Decrease detector voltage: False; Large protein (>70 kDa): False; Ion Source: Source name: TurboIonSpray; Curtain gas: 35 psi; Ion source gas 1: 60 psi; Ion source gas 2: 60 psi; Temperature: 500 ˚C; Experiment: Scan type: TOF MS; Polarity: Positive; Spray voltage: 5500 V; CAD gas: 7; Time bins to sum: 4; Channel 1-4: True; TOF start mass: 350 Da; TOF stop mass 1500 Da; Accumulation time: 0.25; Declustering potential: 80V; Declustering potential spread: 0 V; Collision energy: 10V; Collision energy spread: 0 V; Override Qjet RF value: False.

**2.7 Plasma stability assay**

A 10% Human serum solution was prepared in DPBS (pH = 7.3). From the peptide stock (1 mM in $H_2O$) peptide was diluted to a final concentration of 60 µM in the 10% serum solution. The mixture was mixed immediately after protein addition and 2 µL (for DuoMYC **5**) or 10 µL (for MonoMYC **4**) aliquotes were mixed with 2 µL or 10 µL of a 20% TFA solution in $H_2O$ (T = 0) to quench the human serum, resulting in protein precipitating, and stored on ice for 30 min. Subsequently, the peptide serum solution was incubated at 37 ˚C. At indicated timepoints 2 µL aliquotes (DuoMYC **5**) or 10 µL (MonoMYC **4**) were mixed with 2 µL or 10 µL of 20% TFA in $H_2O$, respectively. Samples were incubated on ice for 30 min and subsequently, the pellet was diluted 5.5x with addional DPBS to redissolve the precipitated protein pellet. This solution was analyzed according to the high resolution LCMS protocol (Method B). As internal standard (IS), the extracted-ion chromatogram (XIC) from 1211.08 was used belonging to a serum protein present in the solution. After measurement, the XIC belonging to the protein, peak 873.1 for MonoMyc **4** and 786.3 for DuoMYC **5**, and IS were extracted from the total-ion chromatogram (TIC) and the background was removed. Using Graphpad Prism 9, the area under the curve (AUC) from each timepoint was calculated between 2.4 min and 3.6 min to obtain the AUC corresponding to the protein, normalized against the AUC of IS followed by normalization against T = 0 and plotted. Next, a nonlinear regression – One phase decay analysis was performed to obtain $t_{1/2}$ with a plateau constant equal to 0 and $Y_0$ set to 1. The assay was performed in independent duplicates.

**3 Protein synthesis with LCMS spectra**

**3.1 MiniMyc with lysine linker (1)**

Fmoc-Lys(Fmoc)-OH (295 mg, 0.5 mmol, 1 eq.) was dissolved in a HATU solution (0.45M, 0.9 eq.) in DMF (1 mL). Subsequently, DIPEA (200 µL, 2.3 eq.) was added to the amino acid solution. The solution was incubated and shaken for 30 seconds and then added to the Chemmatrix Rink amide resin (50 mg, 41 mmol/g). After 10 min incubation at RT, with occasional shaking, the resin was drained and washed with DMF (3 x 5 mL). Next, the peptidyl resin was washed with 20% piperidine in DMF (5 mL) followed by 5 min incubation of 20% piperidine in DMF (5 mL). The resin was drained by vacuum and washed with DMF (5x 5 mL). The dimeric sequence was completed using the standard ASPPS protocol. The final AA was acetylated and cleaved according to the general protocols.

MiniMyc **1** was purified using a Biotage® Selekt Flash Purification System coupled to a reverse phase Biotage® Sfär C18 D - Duo 100 Å 30 µm 25g column, further revered to as reversed phase flash chromatograpy (RP-FC). The mobile phase solvents used were: solvent A ($H_2O$ with 0.1% TFA) and solvent B (MeCN with 0.1% TFA). The UV detector was set to 214 nm. The following method was applied to the crude protein mixture:

RP-FC method: 10% solvent B over 1 column volumes (CV), followed by a linear gradient of 10% solvent B to 40% solvent B over 25 CV, followed by a linear gradient of 40% solvent B to 70% solvent B over 2 CV. The fractions were analyzed via LC-MS (Method A). Pure fractions were combined and lyophilized. Yield: 14.7 mg, 16%.

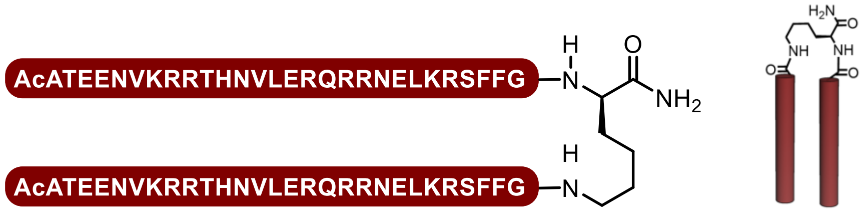


MiniMyc with lysine linker sequence :

(AcATEENVKRRTHNVLERQRRNELKRSFFG)$_2$K

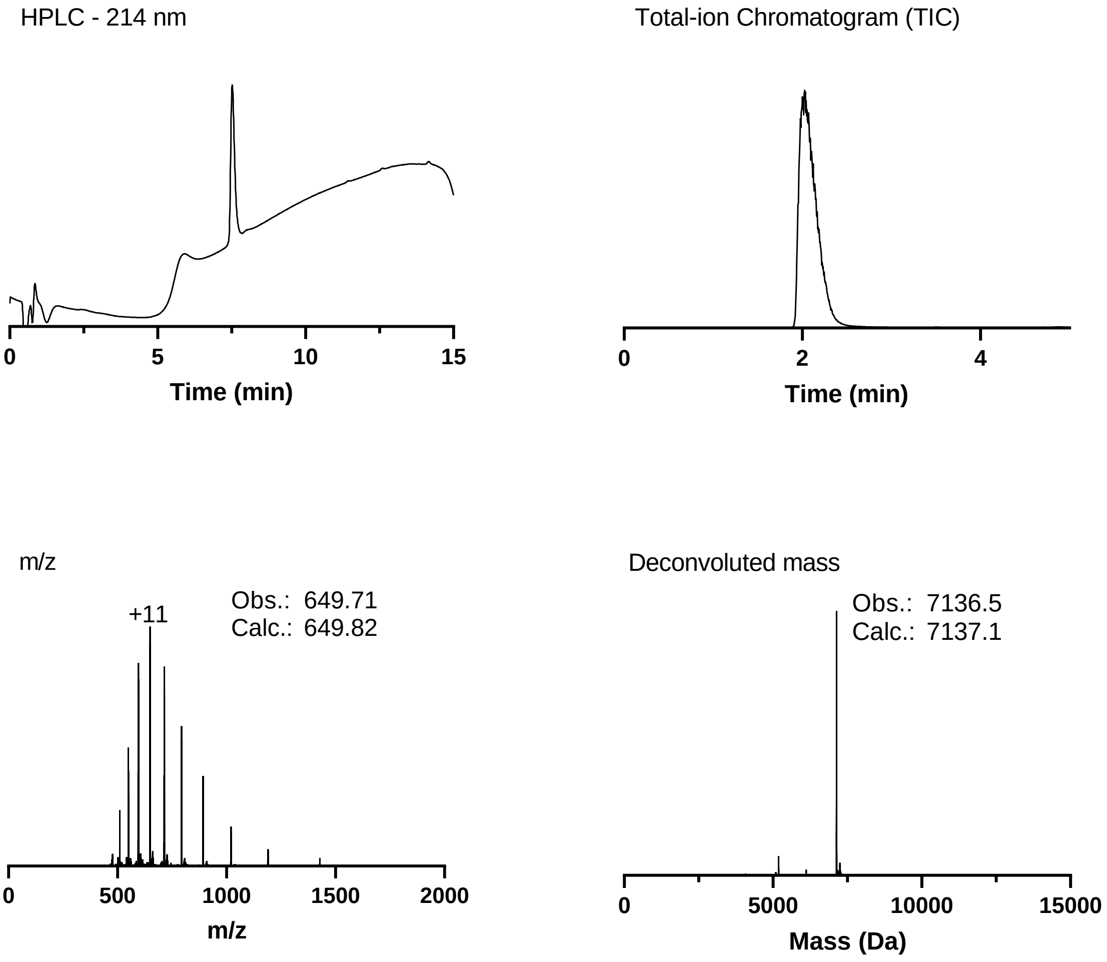


LCMS analytics of MiniMYC **1**. The 214 nm LC chromatogram was obtained using LC-MS method A. The TIC, MS and deconvoluted mass chromatograms were obtained using LC-MS method B.

#### 3.2 MiniMyc monomer (2)

MiniMyc monomer **2** was synthesized, acetylated and cleaved according to the general ASPPS, acetylation and cleavage protocol, respectively.

MiniMyc monomer **2** was purified using a Biotage® Selekt Flash Purification System with a Biotage® Sfär C18 D - Duo 100 Å 30 µm 25g column. The mobile phase solvents used were: solvent A ($H_2O$ with 0.1% TFA) and solvent B (MeCN with 0.1%TFA). The UV detector was set to 214 nm. The following method was applied to the crude protein mixture:

RP-FC method: 10% solvent B over 1 CV, followed by a linear gradient of 10% solvent B to 50% solvent B over 25 CV, followed by a linear gradient of 50% solvent B to 70% solvent B over 2 CV. The obtained fractions were analyzed according to the LC-MS – low resolution general protocol. Pure fractions were combined and stored as lyophilized powders. Yield: 23.4 mg, 16%.

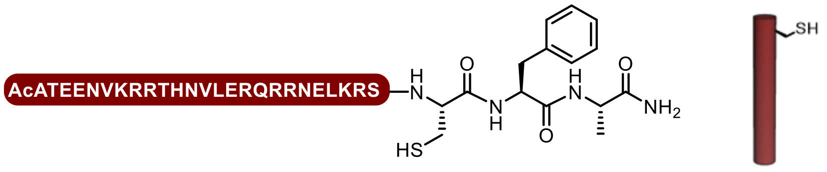


Minimyc monomer sequence :

AcATEENVKRRTHNVLERQRRNELKRSCFA

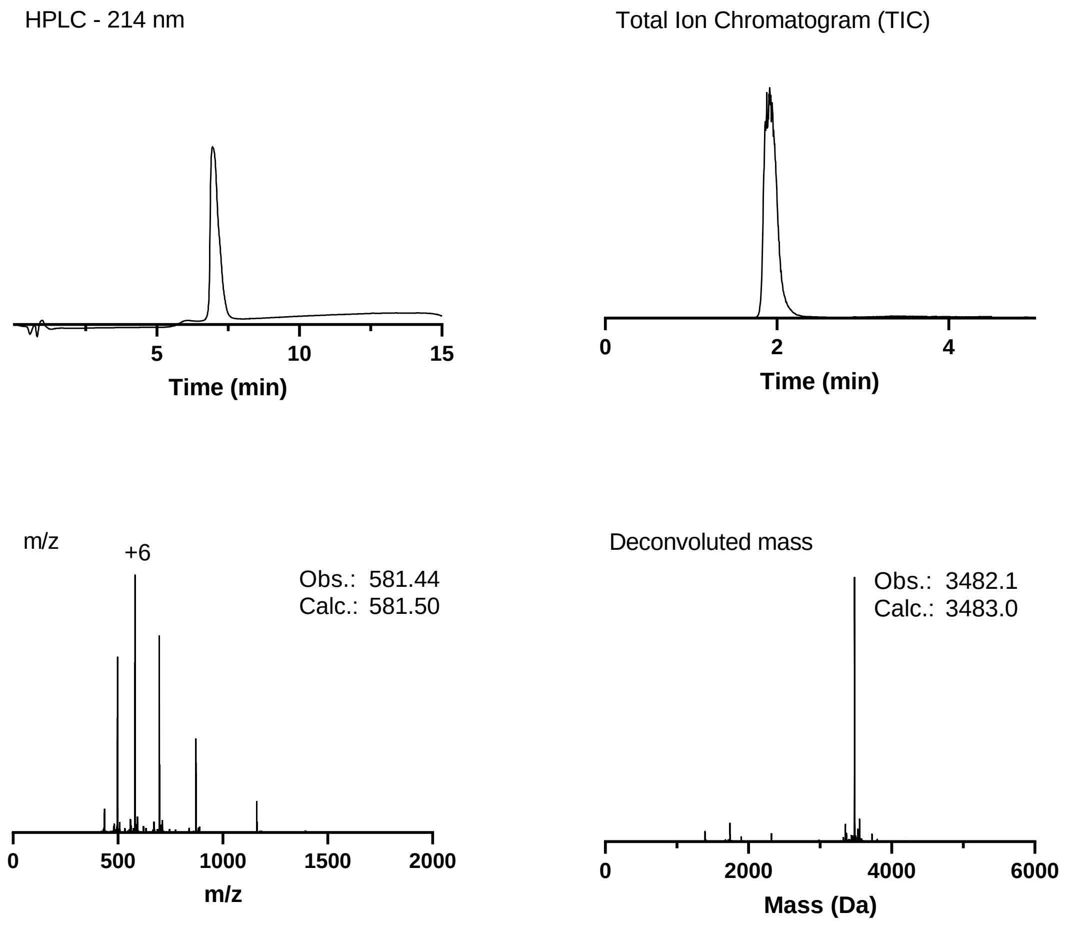


LCMS analytics of MiniMYC monomer **2**. The 214 nm LC chromatogram was obtained using LC-MS method A. The TIC, MS and deconvoluted mass chromatograms were obtained using LC-MS method B.

#### 3.3 MiniMyc with octafluorobiphenyl linker (3)

To prepare peptide **3**, stock solutions were prepared in DMF for decafluorbiphenyl (10 mM), DIPEA (80 mM) and TCEP (20 mM). Peptide **2** (11 mg, 3.16 µmol) was dissolved in 314 µL DMF. From each stock solution 314 µL were added to the peptide solution, resulting in the following final concentrations: peptide (3.16 µmol, 2.5 mM, 1 eq.), decafluorbiphenyl (3.16 µmol, 2.5 mM, 1 eq.), DIPEA (25.3 µmol, 20 mM, 8 eq.) and TCEP (6.32 µmol, 5 mM, 2 eq.) in a final volume of 1.26 mL. The solution was shaken for 24 h after which it was quenched by the addition of 1% TFA in $H_2O$ (10 mL) and lyophilized.

MiniMyc dimer **3** was purified using a Biotage® Selekt Flash Purification System with a Biotage® Sfär C18 D - Duo 100 Å 30 µm 10 g column. The mobile phase solvents used were: solvent A ($H_2O$ with 0.1% TFA) and solvent B (MeCN with 0.1%TFA). The UV detector was set to 214 nm. The following method was applied to the crude protein mixture:

RP-FC method: 10% solvent B over 1 CV, followed by a linear gradient of 10% solvent B to 50% solvent B over 25 CV, followed by a linear gradient of 50% solvent B to 70% solvent B over 2 CV. The obtained fractions were analyzed according to the LC-MS – low resolution general protocol. Pure fractions were combined and stored as lyophilized powders. Yield 3.5 mg, 31%.

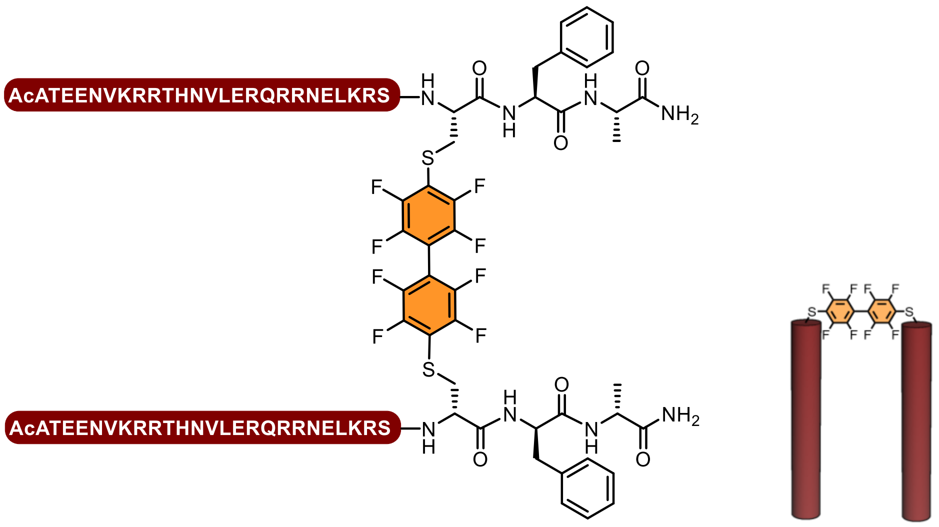


MiniMyc with octafluorobiphenyl linker sequence:

$(\text{AcATEENVKRRTHNVLERQRRNELKRSSCFA})_2$-Linker

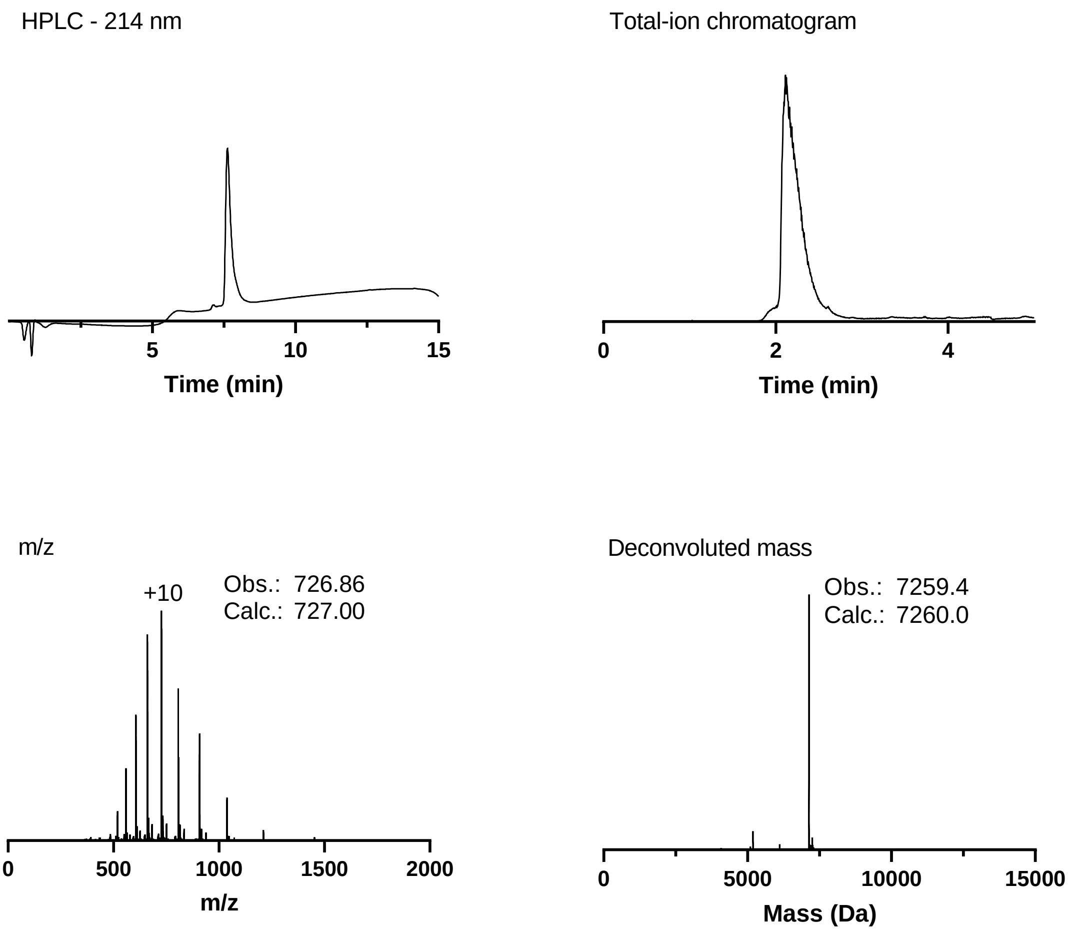


LCMS analytics of MiniMYC **3**. The 214 nm LC chromatogram was obtained using LC-MS method A. The TIC, MS and deconvoluted mass chromatograms were obtained using LC-MS method B.

#### 3.4 MonoMyc (4)

MonoMyc **4** was synthesized according to the general SPPS protocol on a 104 µmol scale. Instead of single couplings, double couplings were performed before removal of the Fmoc protection group at each cycle.

MonoMyc **4** was purified twice. The first purification step was performed using a Biotage® Selekt Flash Purification System with a Biotage® Sfär C18 D - Duo 100 Å 30 µm column 25g. The mobile phase solvents used were: solvent A ($H_2O$ with 0.1% TFA) and solvent B (MeCN with 0.1%TFA). Solvent C (94.9%/5%/0.1% = $H_2O$/ MeCN /TFA) was used to dissolve crude protein mixtures. The UV detector was set to 214 nm. The following method was applied to the crude protein mixture:

RP-FC method: 10% solvent B over 2 CV, followed by 20% solvent B over 3 CV, followed by 25% solvent B over 2 CV, followed by 29% solvent B over 3 CV, followed by 32% solvent B over 4 CV, followed by 40% solvent B over 3 CV, followed by 98% solvent B over 5CV. The obtained semipure fractions where analyzed using a Sciex X500b QTOF LC/MS (see LC-MS high resolution protocol). Fractions containing product were combined and lyophilized.

The second purification was performed according to the general prep-HPLC protocol. Yield: 3.6 mg, 0.5%.

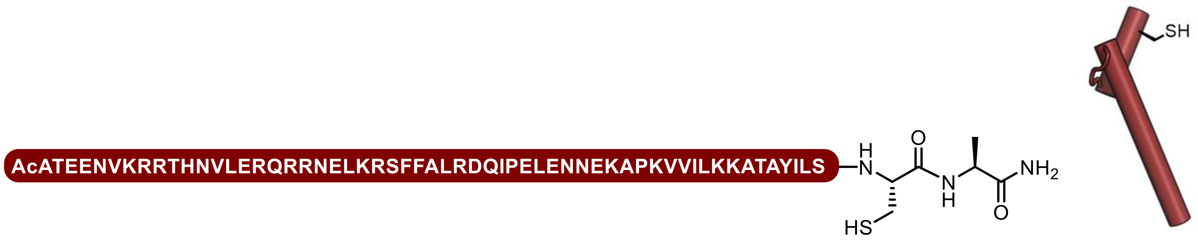


MonoMyc long sequence:

AcATEENVKRRTHNVLERQRRNELKRSFFALRDQIPELENNEKAPKVVILKKATAYILSCA

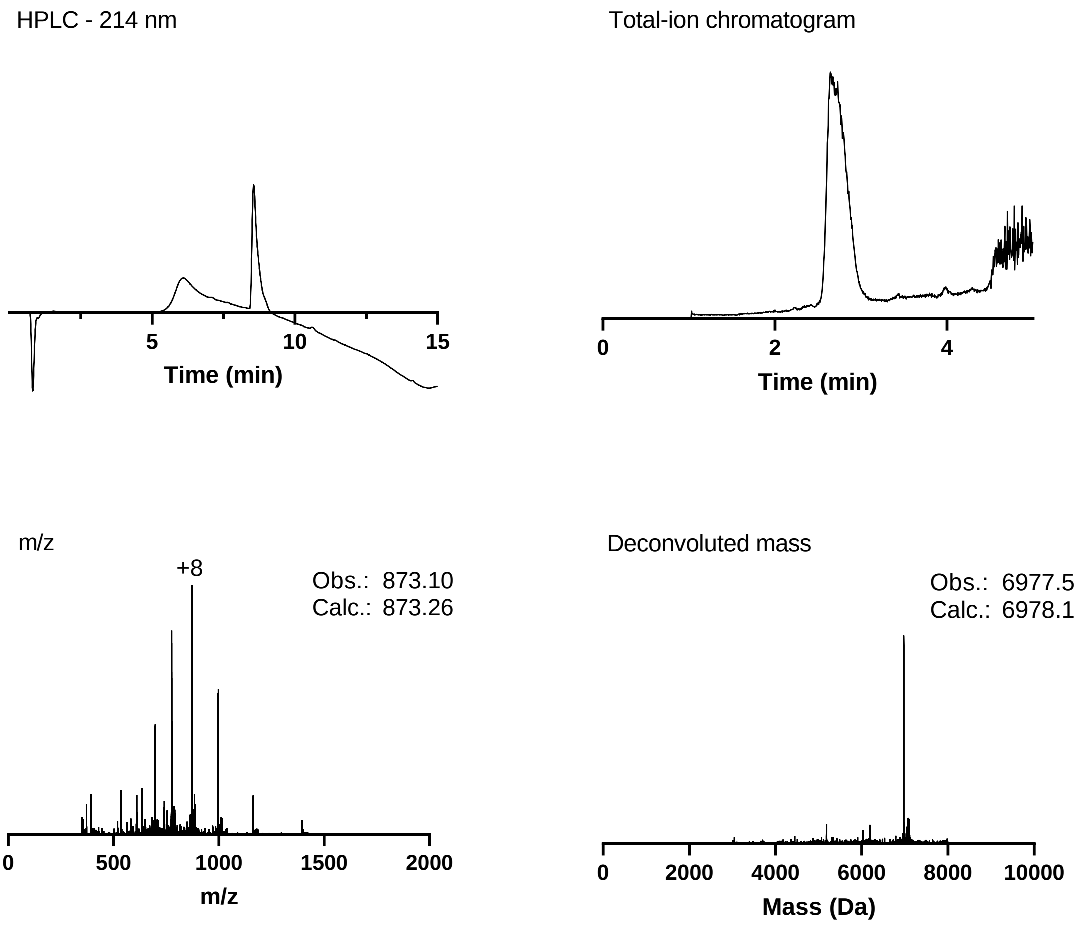


LCMS analytics of MonoMyc **4**. The 214 nm LC chromatogram was obtained using LC-MS method A. The TIC, MS and deconvoluted mass chromatograms were obtained using LC-MS method B.

#### 3.5 DuoMyc with 4,4'-dimethyl-1,1'-biphenyl linker (5)

Peptide **4** (6 mg, 0.86 µmol, 1 eq.) was dissolved in $H_2O$ (86 µL). A stock solution was prepared for 4,4′-Bis(bromomethyl)biphenyl in MeCN (4 mM). The peptide **4** solution was equally divided over 12 eppendorf tubes (7.2 µL). To each eppendorf tube 46.6 µL of of $NH_4HCO_3$ buffer (100 mM, pH = 8) and 17.9 µL of the 4,4′-Bis(bromomethyl)biphenyl stock were added, resulting in a final volume of 71.7 µL and the following concentrations: peptide **4** (0.07 µmol, 1 mM, 1 eq.) and 4,4′-

Bis(bromomethyl)biphenyl (0.07 µmol, 1 mM, 1 eq.). The tubes were shaken for 3h and subsequently lyophilized. The lyophilized reactions where dissolved in a solution of solvent A (94.9%/5%/0.1% = $H_2O$/ MeCN /TFA) and combined. After confirmation by low resolution LC-MS the dimer was purified according to the prep-HPLC protocol as described in 2.4. Yield: 1.4 mg, 23%

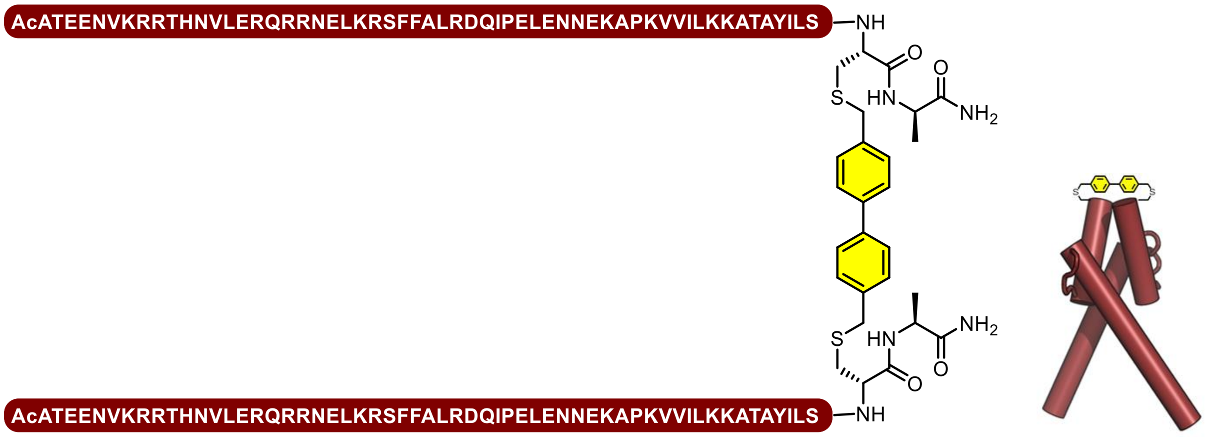


DuoMyc with 4,4'-dimethyl-1,1'-biphenyl linker sequence:

(AcATEENVKRRTHNVLERQRRNELKRSFFALRDQIPELENNEKAPKVVILKKATAYILSCA)$_2$-linker

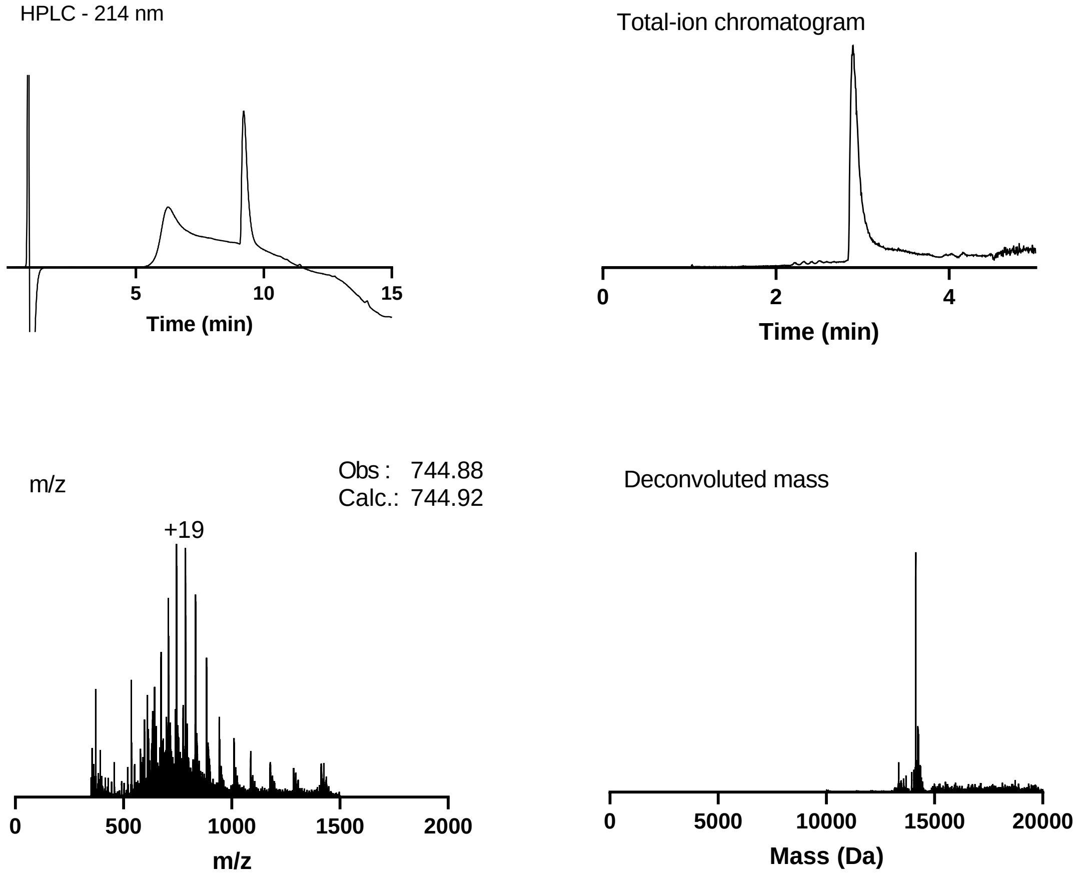


LCMS analytics of DuoMYC **5**. The 214 nm LC chromatogram was obtained using LC-MS method A. The TIC, MS and deconvoluted mass chromatograms were obtained using LC-MS method B.

#### 3.6 MonoMyc (6)

MonoMyc short **6** was synthesized according to the general ASPPS, acetylation and cleavage protocols on a 41 µmol scale. MonoMyc short **6** was purified using a Biotage® Selekt Flash Purification System with a Biotage® Sfär C18 D - Duo 100 Å 30 µm column 25g. The mobile phase solvents used were: solvent A ($H_2O$ with 0.1% TFA) and solvent B (MeCN with 0.1% TFA). The UV detector was set to 214 nm. The following method was applied to the crude protein mixture:

RP-FC method: 20% solvent B over 2 CV, followed by a linear gradient of 20% solvent B to 30% for 3 CV, followed by a linear gradient of 30% solvent B to 40% over 22 CV, followed by 100% solvent B over 3 CV. The obtained semipure fractions were analyzed using the low resulution LCMS protocol. Fractions containing product were combined and lyophilized. Yield: 17 mg, 6%.

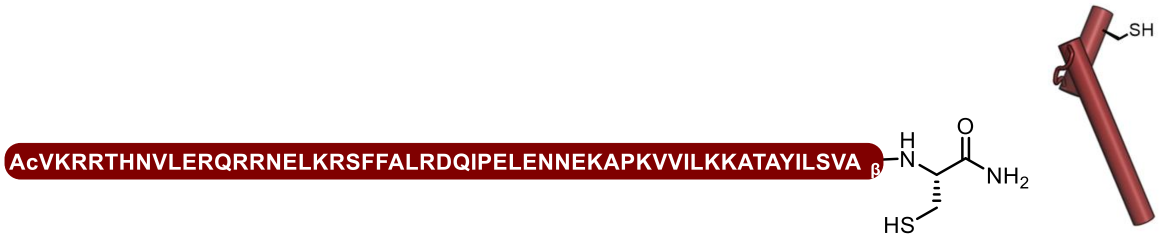


MonoMyc sequence:

AcVKRRTHNVLERQRRNELKRSFFALRDQIPELENNEKAPKVVILKKATAYILSVA$_{\beta}$C

HPLC - 214 nm

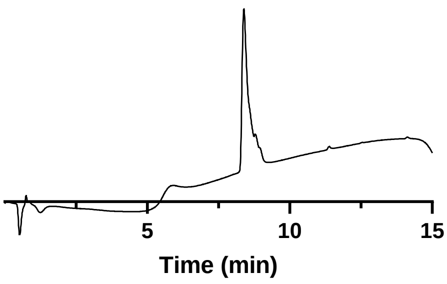


Total-ion chromatogram

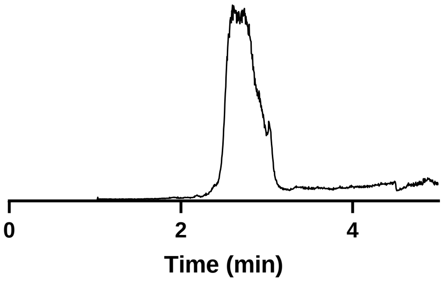


m/z

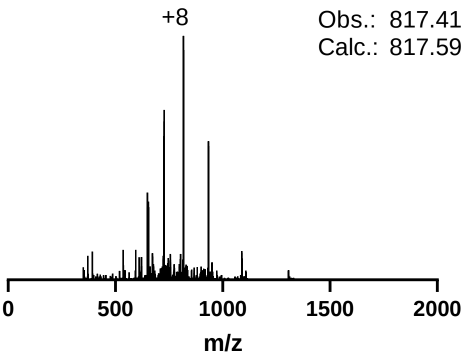


Deconvoluted mass

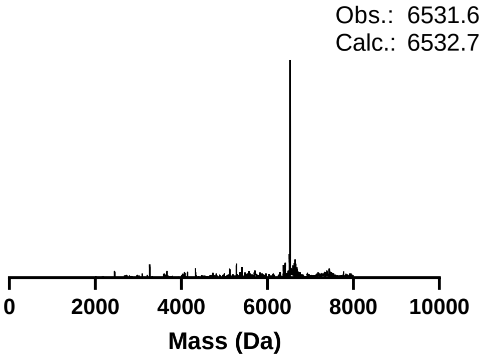

LCMS analytics of MonoMYC **6**. The 214nm LC chromatogram was obtained using LC-MS method A. The TIC, MS and deconvoluted mass chromatograms were obtained using LC-MS method B.

#### 3.7 DuoMyc with octafluorobiphenyl linker (7)

MonoMyc short **6** (4.24 mg, 0.64 µmol, 1 eq.) was disolved in DMF (58 µL). The following stock solutions were prepared in DMF: DIPEA (0.5M), decalfluorobiphenyl (0.01M) and TCEP (0.1M). To the MonoMyc short **6** solution was added from each stock solution: 13 µL of DIPEA (6.4 µmol, 10 eq., 60 mM), 19.5 µL TCEP (1.2 µmol, 0.3 eq., 17.7 mM) and 19.5 µL decafluorbiphenyl (0.19 µmol, 3 eq., 1.77 mM) resulting in a final peptide concentration of 6 mM. The reaction was shaken for 3 days at 37°C. When precipitation occurred during the reaction, the reaction mixture was sonicated, vortexed and placed back in the shaker. Yield: 1.28 mg, 12%.

DuoMyc **7** was purified using a Biotage® Selekt Flash Purification System with a Biotage® Sfär C18 D - Duo 100 Å 30 µm column 10g. The mobile phase solvents used were: solvent A ($H_2O$ with 0.1% TFA, v/v) and solvent B (MeCN with 0.1%TFA, v/v). The UV detector was set to 214 nm. The following method was applied to the crude protein mixture:

HPFC method: 20% solvent B over 2 column volumes (CV), followed by a linear gradient of 20% solvent B to 30% for 3 CV, followed by a linear gradient of 30% solvent B to 40% over 22 CV, followed by 100% solvent B over 3 CV. The obtained semipure fractions were analyzed using the low resolution LCMS protocol. Fractions containing product were combined and lyophilized. Yield: 1.28 mg, 12%.

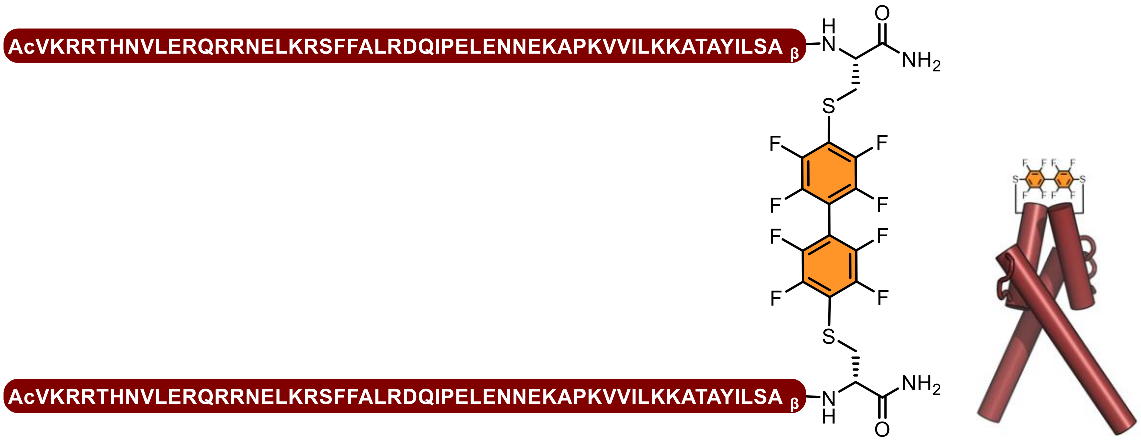


DuoMyc with octafluorobiphenyl linker sequence:

(AcVKRRTHNVLERQRRNELKRSFFALRDQIPELENNEKAPKVVILKKATAYILSA$_\beta$C)$_2$-linker

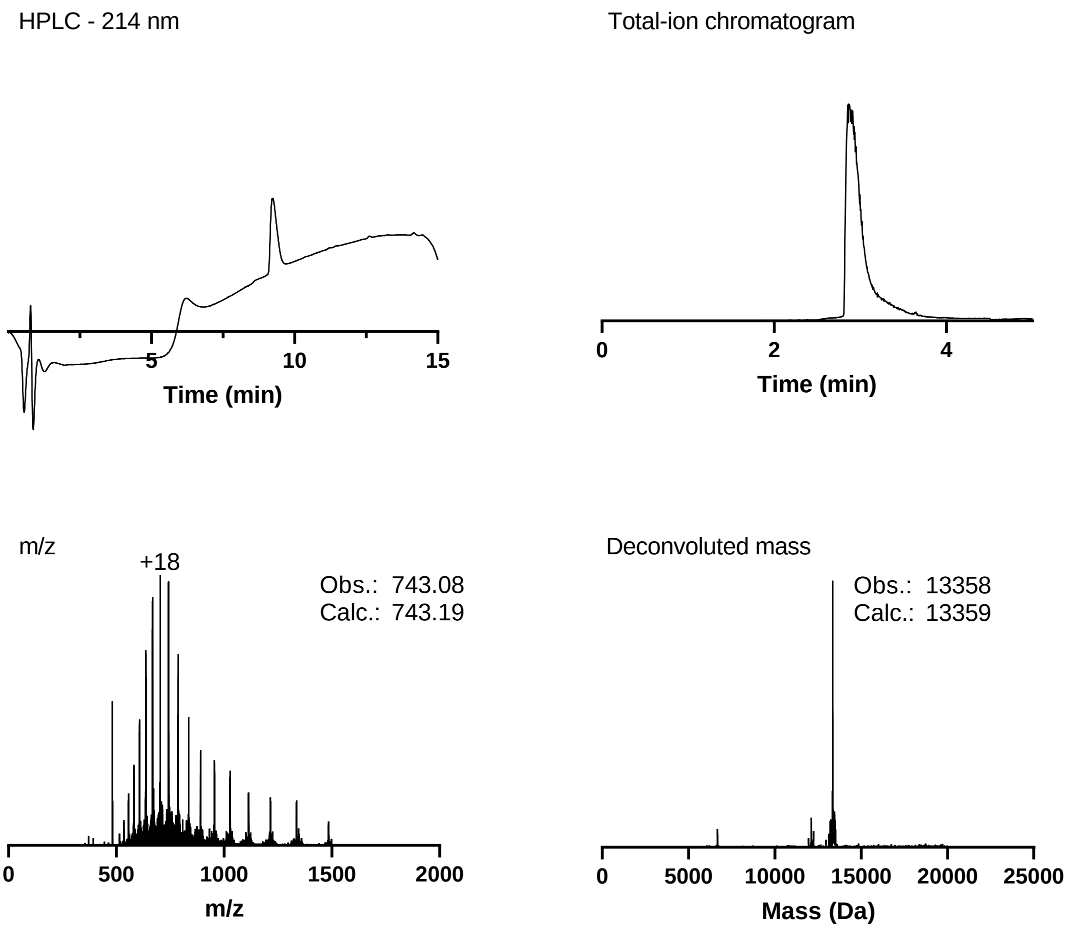


LCMS analytics of DuoMYC **7** The 214 nm LC chromatogram was obtained using LC-MS method A. The TIC, MS and deconvoluted mass chromatograms were obtained using LC-MS method B.

### 4. Biochemistry

#### 4.1. Overview of buffers used

| | |
|---|---|
| Bacterial cell lysis buffer | 20mM Tris-HCl, pH 8, 0.5 M NaCl, 3mM $MgCl_2$, 0.05% DNAse,cOmplete EDTA free protease inhibitor (1 tablet freshly added in 50 mL buffer) |
| EK cleavage buffer | 200 mM Tris-HCl, pH 7.4, 0.5 M NaCl, 20 mM $CaCl_2$ |
| EK dilution buffer | 20 mM Tris-HCl, pH 7.4, 200 mM NaCl, 2 mM $CaCl_2$, 50% glycerol |
| Affinity wash buffer | 20 mM Tris-HCl, pH 8, 0.5 M NaCl, 10mM imidazole |
| Affinity elution buffer | 20 mM Tris-HCl, pH 8, 0.5 M NaCl, 500mM imidazole |
| EMSA binding buffer | 20 mM HEPES, pH 8.0, 150 mM NaCl, 5% glycerol, 1 mM EDTA, 2 mM $MgCl_2$, 0.5 mg/mL of BSA, 1 mM DTT and 0.05% NP-40 |
| EMSA running buffer | 0.5x TBE, 50 mM Tris borate, pH 8.3, 1 mM EDTA |

#### 4.2. Expression of Omomyc in E.Coli

Plasmid pET-30 with Omomyc was gratefully received from the research group of Prof. Dr. Eilers at University of Wuerzburg (see annex for plasmid sequence) and sequenced before use.

Plasmids were transformed into competent ArcticXpress DE3 RIL cells (Agilent) by heat shock. 2 µL of 10% β-mercaptoethanol were mixed with 100 µL of competent cell suspension thawed on ice and incubated for 10 minutes on ice. Next, 25 ng of plasmid DNA were added and the cells incubated for another 30 minutes on ice. The cells were then heat-shocked in a water bath for 20 seconds at 42 °C and subsequently incubated on ice for 2 minutes followed by the addition of 0.9 mL SOC media and incubation at 37 °C, 250 rpm for 1 h. Cells were pelleted by centrifugation, 0.9 mL of the supernatant decanted and the pellet resuspended in the remaining 100 µL of media. Subsequently, cells were plated for selection on LB agar with kanamycin and gentamicin and incubated at 37 °C overnight.

Single colonies were picked from the plate and cultured overnight at 37 °C and 200 rpm in 100 mL of LB media containing kanamycin and gentamicin. Next, 3x2 L of LB media containing kanamycin and gentamicin in 5 L Erlenmeyer flasks were inoculated with 25 mL of preculture and grown at 37 °C, 200 rpm until an OD of 0.8 was reached. The temperature was then set to 14 °C and protein expression induced by addition of 100 µM of IPTG. Protein expression was conducted overnight for 18 h after which the cells were harvested by centrifugation (5,000g, 4 °C, 12 min.), the pellet resuspended in lysis buffer (20mM Tris-HCl, pH 8, 0.5 M NaCl, 3mM $MgCl_2$, 0.05% DNAse,cOmplete EDTA free protease inhibitor) and the cells lysed by pressure lysis. Cell debris was removed by ultracentrifugation (35,000 rpm, 4°C, 45 min.).

#### 4.3. Purification of his-tagged proteins

The supernatant after ultracentrifugation was purified using an ÄKTA start protein purification system equipped with a 5 mL HisTrap HP His tag protein purification column (Cytiva). After washing out unbound protein with wash buffer (20 mM Tris-HCl pH 8, 0.5 M NaCl, 10 mM imidazole) the protein was eluted using a gradient of 10 mM to 500 mM imidazole in the same buffer over 40-50 column volumes. Fractions with protein were analyzed for protein content and purity by SDS-PAGE.

#### 4.4. General protocol for SDS-PAGE

Samples were mixed with the appropriate amount of 4x LB buffer containing β-mercaptoethanol and heated to 95 °C for 5 minutes to denature proteins. Samples were then loaded onto acrylamide gels of the desired percentage (made using acrylamide 37.5:1 acrylamide/Bis) and ran at room temperature at 150V for one hour. If desired, gels were stained for at least 20 minutes with coomassie blue staining solution followed by at least three rounds of destaining with destaining solution (50% MeOH, 40 % $H_2O$, 10% AcOH). Stained gels were scanned on a Bio-Rad ChemiDoc MP machine.

**4.5. General protocol for buffer exchange**

The combined fractions obtained from Ni-column purification were incubated with TCEP (500 µM-1mM) to break any possible formed disulfide bonds. The buffer was then exchanged by subjecting the protein to column chromatography on a Biotage® Selekt Flash Purification System equipped with a Biotage® Sfär C18 D - Duo 25g or 50g column and a stepwise gradient of 0 % MeCN in water, 50 to 100% MeCN water). The combined fractions containing the protein were lyophilized, yielding the His-tagged or final protein as TFA salt.

**4.6. His-tag cleavage by enterokinase**

Omomyc was dissolved at 2 mg/mL in EK cleavage buffer (200 mM Tris-HCl, pH 7.4, 0.5 M NaCl, 20 mM $CaCl_2$) and after addition of 10 u/mL enterokinase (previously diluted in dilution buffer, 20 mM Tris-HCl, pH 7.4, 200 mM NaCl, 2 mM $CaCl_2$, 50% glycerol), the protein was incubated overnight. Cleaved Omomyc was purified purified using an ÄKTA start protein purification system equipped with a 5 mL HisTrap HP His tag protein purification column with the protein being eluted in the flowthrough. Fractions were checked fro protein content and purity using SDS-PAGE and the buffer was exchanged using the general protol for buffer exchange.

**4.7. Electromobility shift assay (EMSA)**

For EMSAs, 14 µL (15 µL for no protein control) water were mixed with 1 µL of 20x FAM-labelled DNA construct (IRD700-ACC CCA C**CA CGT G**GT GCC T), 1 µL of 20x protein in water and 4 µL of 5x EMSA buffer (final buffer concentration: 20 mM HEPES, pH 8.0, 150 mM NaCl, 5% glycerol, 1 mM EDTA, 2 mM $MgCl_2$, 0.5 mg/mL of BSA, 1 mM DTT and 0.05% NP-40). The samples were incubated for 30 minutes at room temperature, placed on ice and incubated for another 15 minutes after which 15 µL of the samples were loaded onto a 10% acrylamide TBE gel which was pre run before for 1h at 75V at 4 °C in 0.5x TBE. Samples were run for 20 minutes at 120 V followed by 40 minutes at 100 V at 4 °C in 0.5x TBE and subsequently scanned on on a Bio-Rad ChemiDoc MP machine.

**4.8. Cell culture**

Cells were cultured at 37°C in 5% $CO_2$ atmosphere. Cell lines were cultured in ATCC recommended media and split twice a week before confluency was reached.

**4.9. Reporter gene assay**

The reporter gene assay was performed using Cignal reporter assay (CCS-012L, Qiagen) according to the manufacturer's protocol with slightly prolonged incubation times. In brief, HEK293T cells were harvested and resuspended in OptiMEM media containing 5% FBS and 1% non essential amino acids. 40,000 cells were seeded per well in a 96-well plate and transfection cocktail of either cignal reporter or positive or negative control reporter along with attractene transfection reagent was added and the cells incubated overnight. Next, media was changed to assay media (OptiMEM, 0.5% FBS, 1% NEAA, Pen/Strep) and the cells were incubated for 8 hours after which the media was replaced by 75 µL assay media containing the different proteins/control compounds at the required concentration and the cells were incubated for 16-24 hours.

Luciferase assay was then performed using a luciferase assay kit (E2940, Promega). Cells were lysed by addition of 75 µL of DualGlo Luciferase assay reagent and incubated for 15 minutes after which Luciferase luminescence was measured in on a Bio-Rad ChemiDoc MP machine. Subsequently, 75 µL of DualGlo Stop & Glo reagent were added and the Renilla luciferase luminescence measured after 15 minutes of incubation time. Signal was normalized against cell number by calculating the ratio of firefly and renilla luminescence and eventually these signals were normalized against the untreated control. All experiments were done at least in technical duplicates.

**5. Molecular Dynamic simulations**

The pdb structure of omoMYC bound to DNA was retrieved from the PDB (id=5I50) and subsequently prepared for MD using Maestro's 2022-3 protein preparation wizard. Thereafter, a system was build using the desmond system builder, MD simulations were performed using the OPLS4 forcefield under the NPgT ensemble at a temperature of 300K using 2fs timesteps. Simulations were performed in triplicate using a different random seed for the initial velocities, and the resulting 500ns trajectories were subsequently analyzed using the desmond simulation event analysis. All structural images were generated using PyMOL

### 6. Supplementary figures

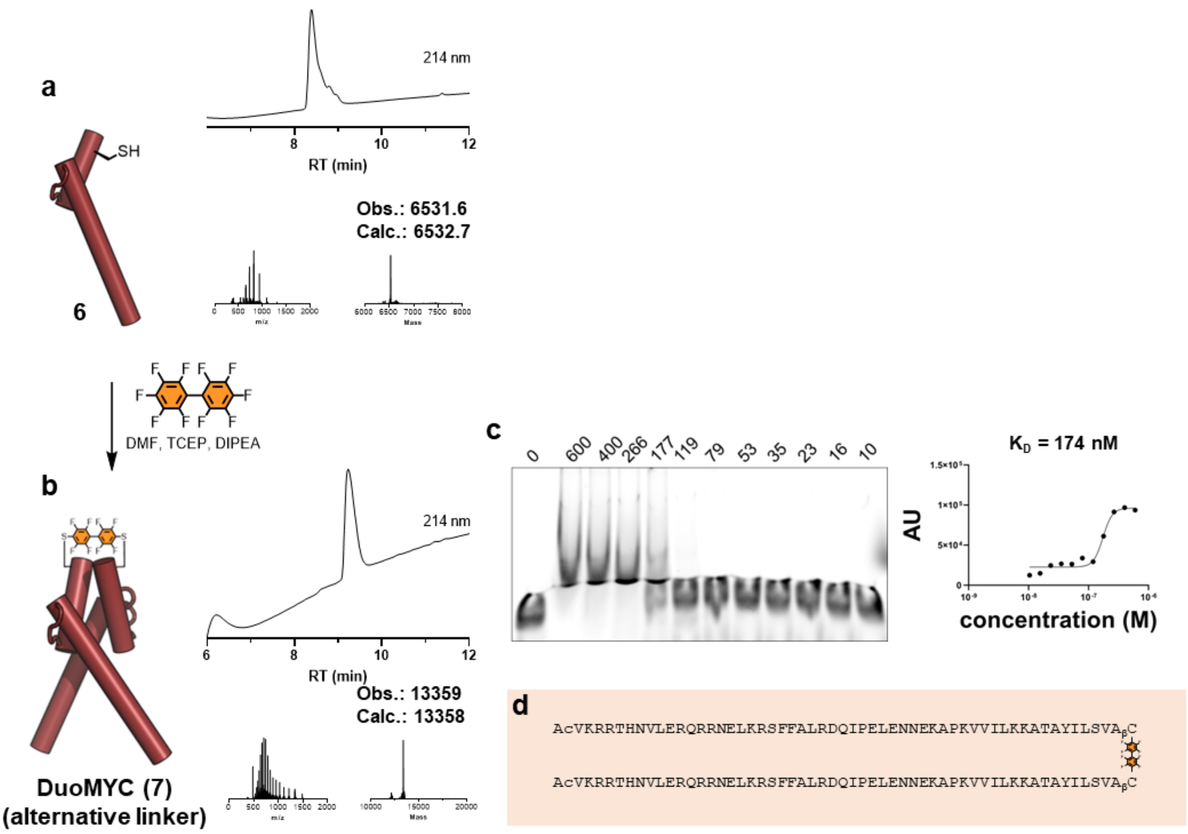


**Figure S1.** DuoMyc (**7**) with alternative (octafluorophenyl) linker binds E-Box with nanomolar affinity. (a) SPPS of monomeric variant **6** with LCMS analysis. b) Dimerization of **6** with decafluorobiphenyl, leading to DuoMYC (**7**). c) DuoMYC (**7**) binds to E-Box with a $K_D$ = 174 nM. d) Sequence of DuoMYC (7). General SPPS coupling conditions: peptidyl resin incubated with Fmoc-AA-OH (10 eq.), HATU (9 eq.) and DIPEA (29 eq.) in DMF for 8 minutes at 70 $^{o}$C. Fmoc was removed with 20% piperidine + 2% formic acid in DMF (4 minutes at 70 $^{o}$C). General EMSA protocol: Fluorescently-labelled dsDNA construct (IRD700-ACCCCACCACGTGGTGCCT, final concentration 4 nM) was preincubated with protein in EMSA buffer (20 mM HEPES, pH 8.0, 150 mM NaCl, 5% glycerol, 1 mM EDTA, 2 mM)

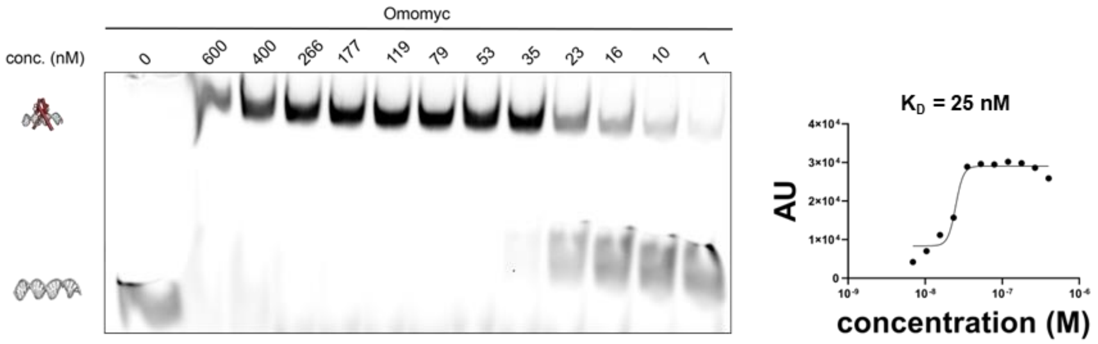


**Figure S2.** Omomyc binds to E-box DNA with an affinity of 25 nM. General EMSA protocol: Fluorescently-labelled dsDNA construct (IRD700-ACCCCACCACGTGGTGCCT, final concentration 4 nM) was preincubated with protein in EMSA buffer (20 mM HEPES, pH 8.0, 150 mM NaCl, 5% glycerol, 1 mM EDTA, 2 mM)

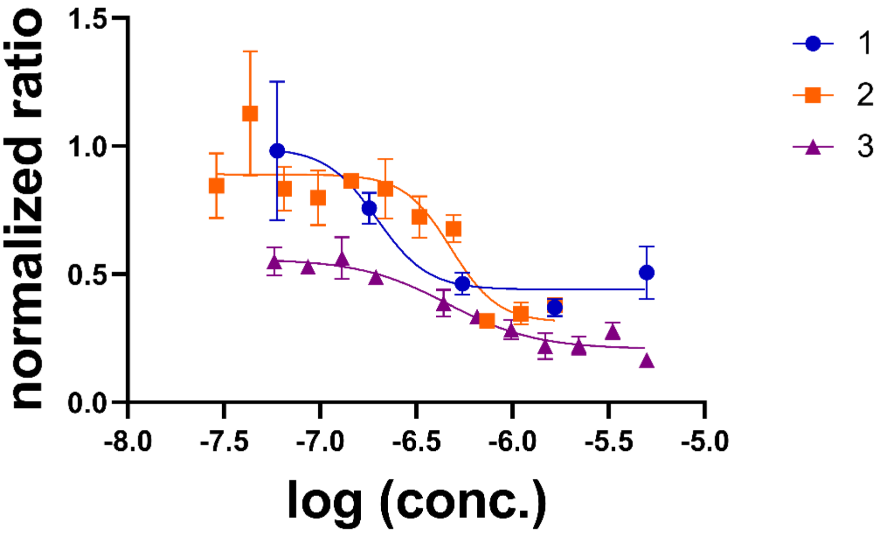


**Figure S3.** Three independant experiments to determine the EC50 of DuoMyc (5) in a reporter gene assay. Curves were fitted to logarithmized data and results indicate EC50s in the nanomolar range. $EC_{50}$(1) = 198 nM, $EC_{50}$(2) = 480 nM, $EC_{50}$(3) = 464 nM.

### Images of uncut gel files

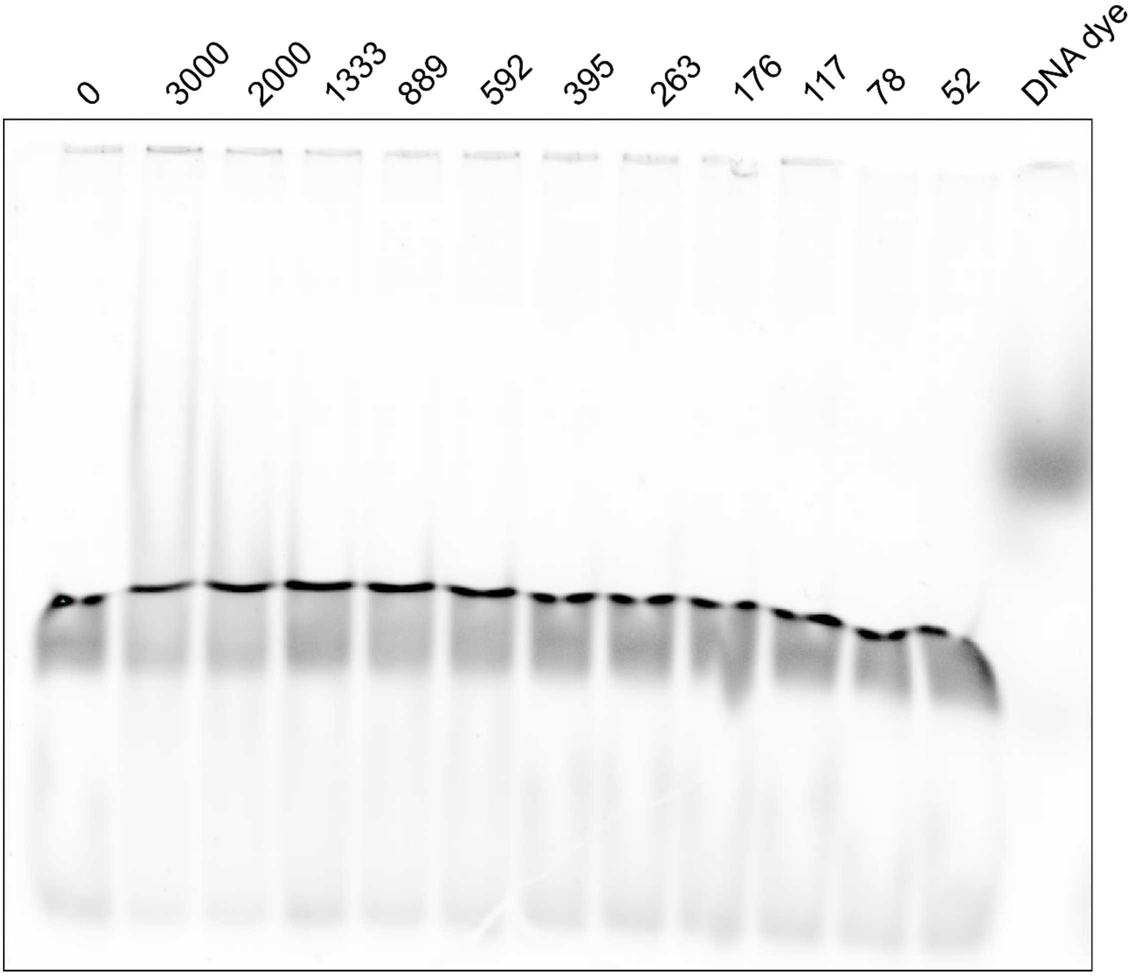


**Figure S4.** Uncut EMSA gel image of miniMyc (**1**), as shown in the main text in figure 2b.

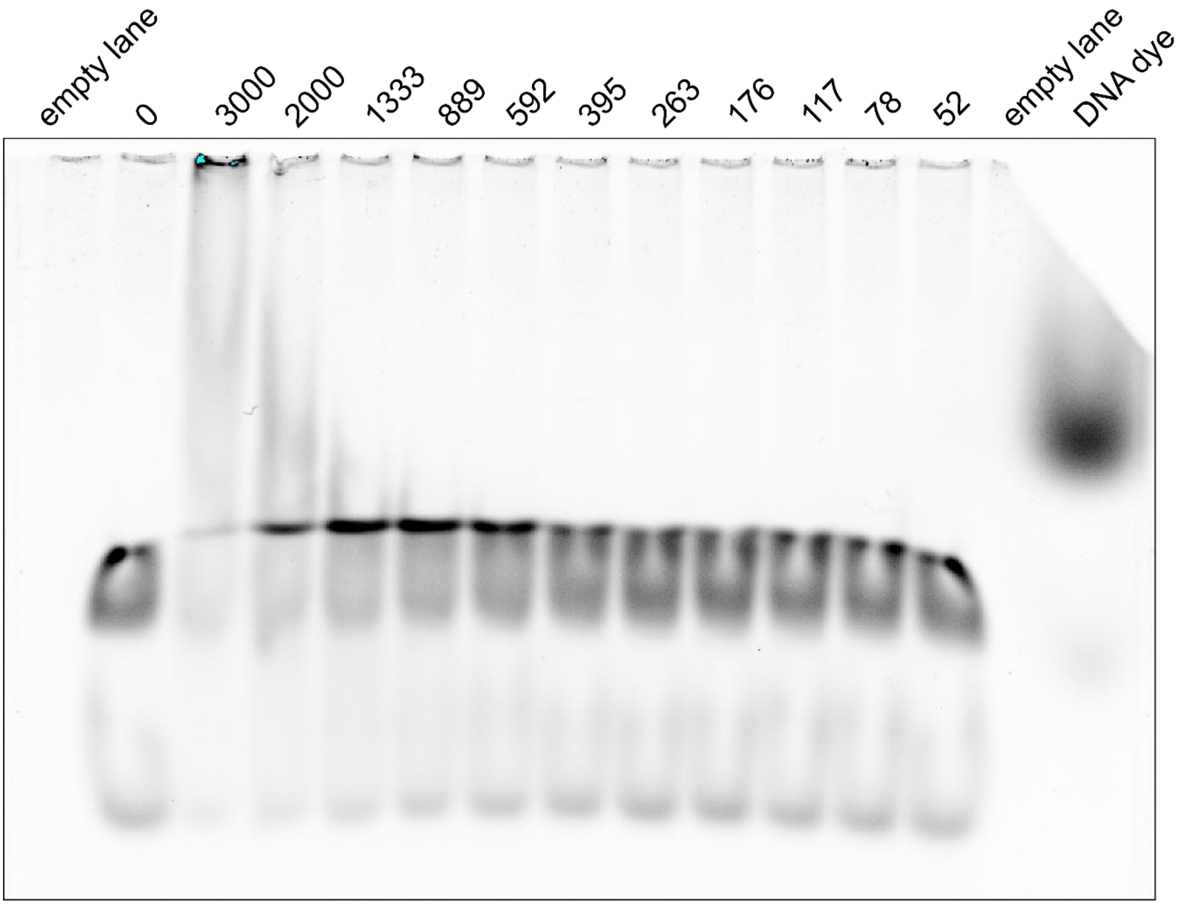


**Figure S5.** Uncut EMSA gel image of miniMyc (**3**), as shown in the main text in figure 2d.

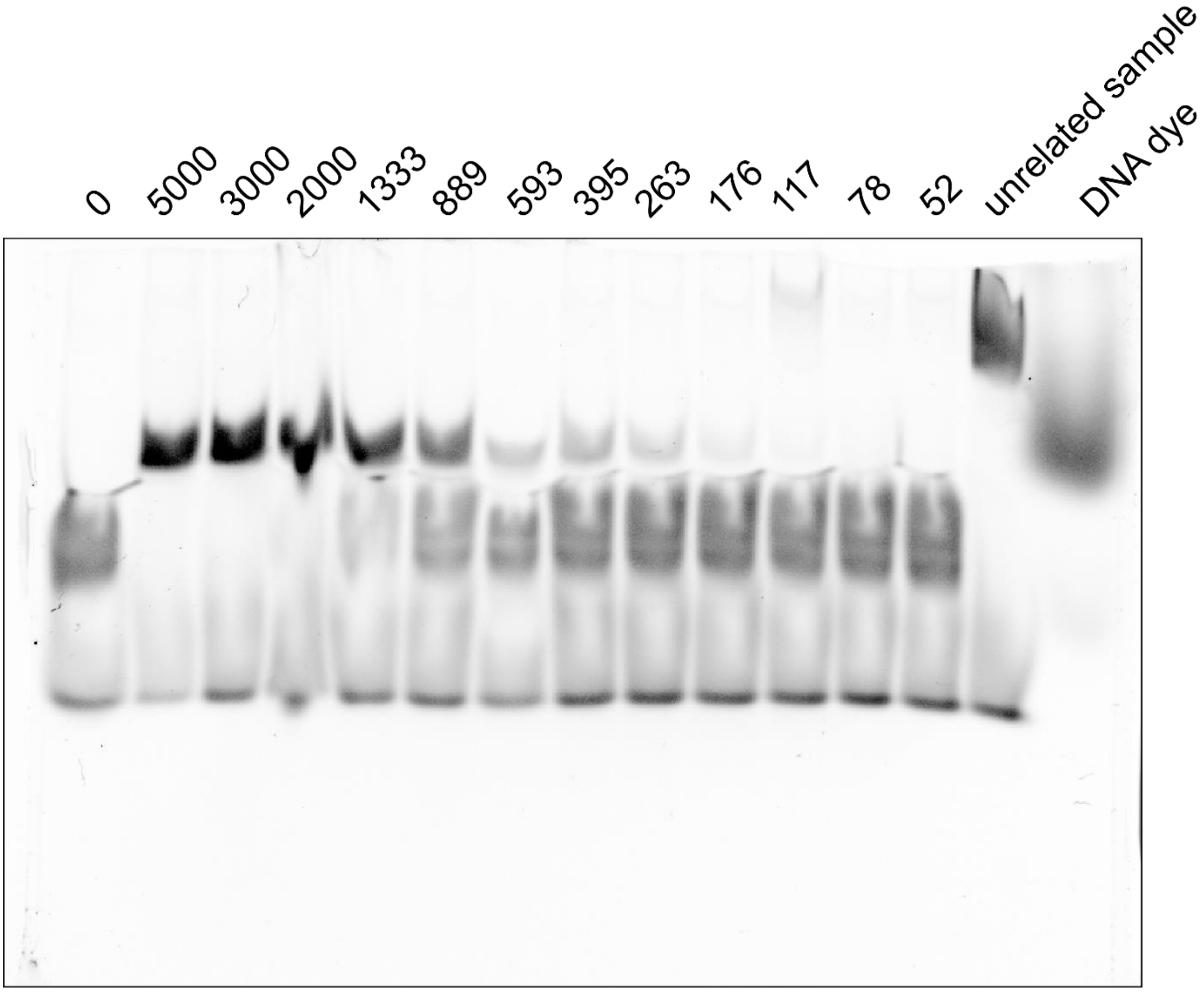

**Figure S6.** Uncut EMSA gel image of DuoMyc monomer (**4**) as shown in the main text in figure 2f.

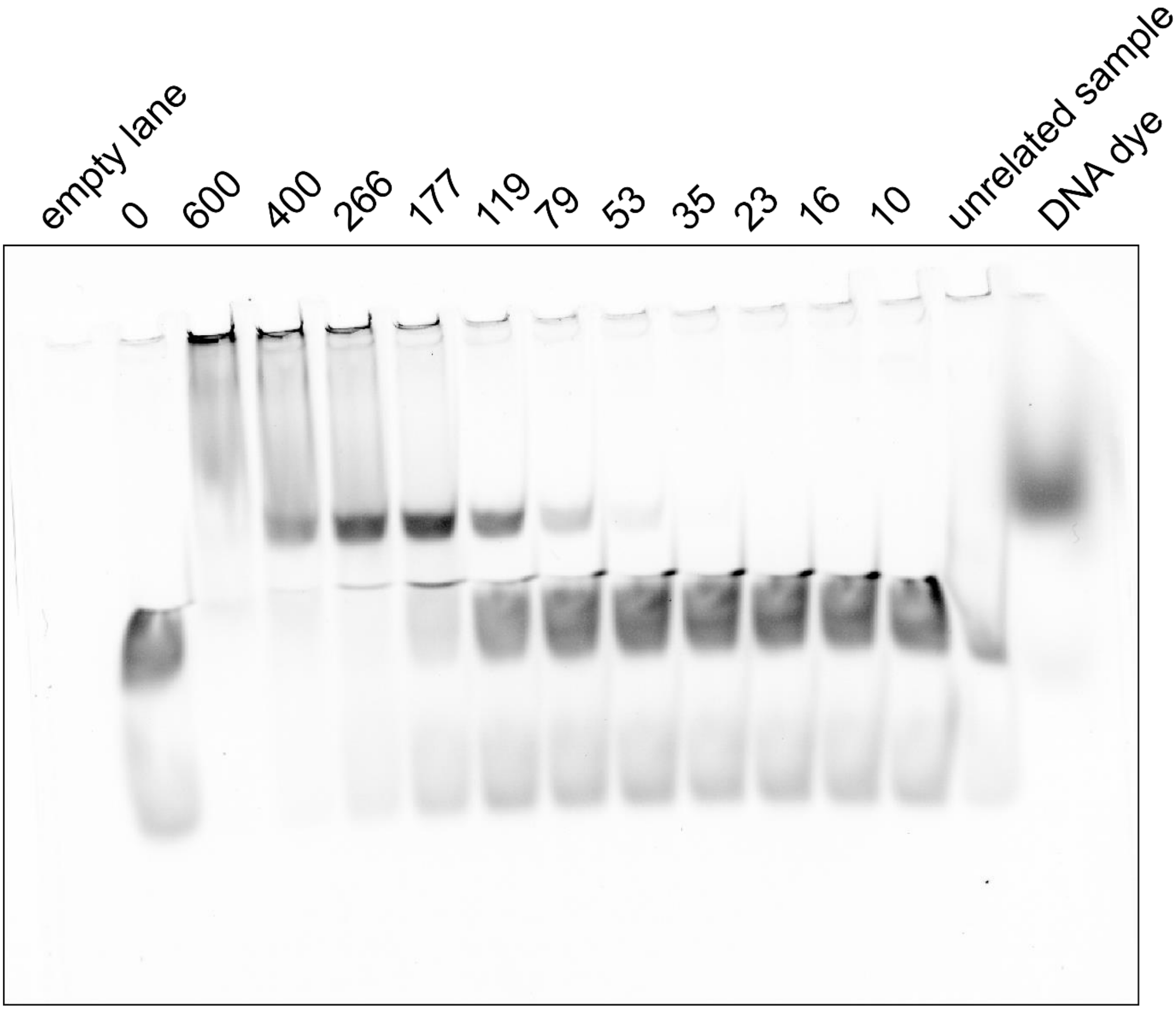

**Figure S7.** Uncut EMSA gel image of DuoMyc (**5**), as shown in the main text in figure 2h.

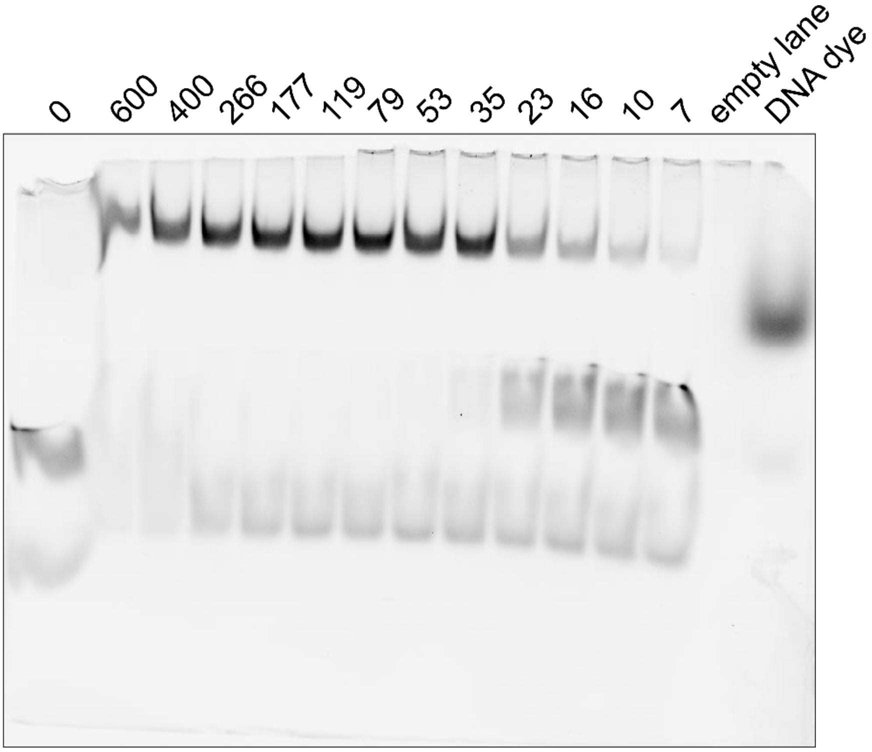


**Figure S8.** Uncut EMSA gel image of Omomyc as shown in figure S2.

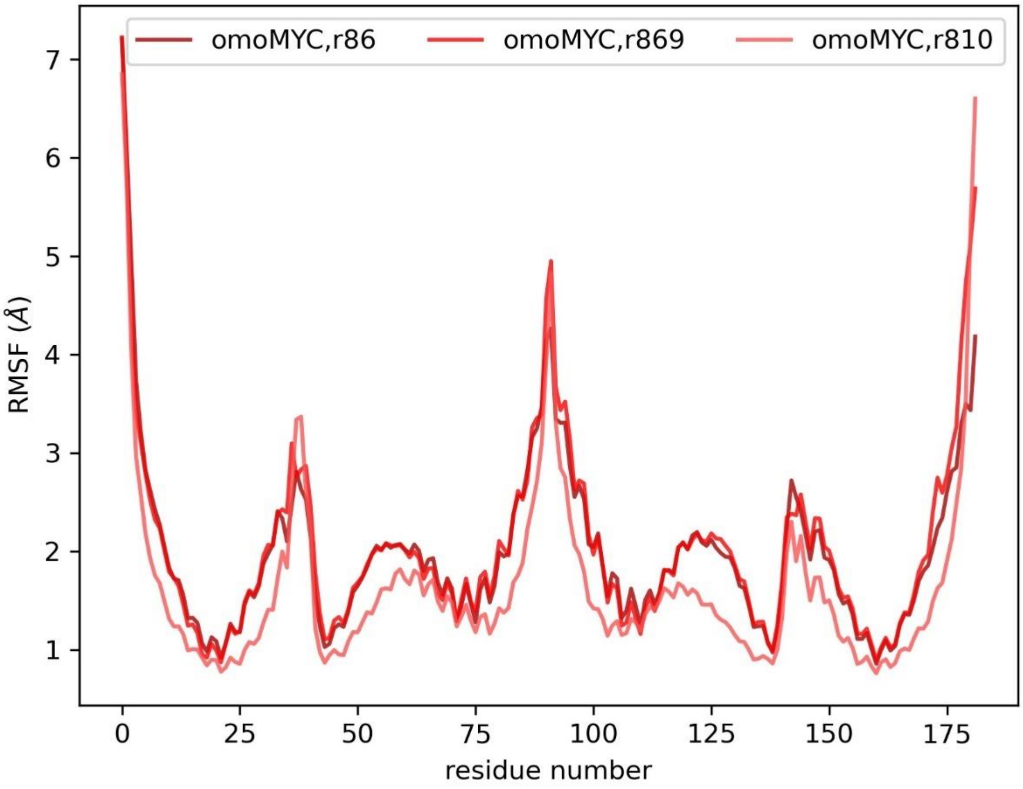


**Figure S9**. RSMF plot of OmoMYC (500 ns MD simulation).

**Plasmide and protein sequences**
**pET-30 a with DuoMyc**

TCAGAGGTTTTCACCGTCATCACCGAAACGCGCGAGGCAGCTGCGGTAAAGCTCATCAGCGTGGTCGT
GAAGCGATTCACAGATGTCTGCCTGTTCATCCGCGTCCAGCTCGTTGAGTTTCTCCAGAAGCGTTAAT
GTCTGGCTTCTGATAAAGCGGGCCATGTTAAGGGCGGTTTTTTCCTGTTTGGTCACTGATGCCTCCGT
GTAAGGGGGATTTCTGTTCATGGGGGTAATGATACCGATGAAACGAGAGAGGATGCTCACGATACGGG
TTACTGATGATGAACATGCCCGGTTACTGGAACGTTGTGAGGGTAAACAACTGGCGGTATGGATGCGG
CGGGACCAGAGAAAAATCACTCAGGGTCAATGCCAGCGCTTCGTTAATACAGATGTAGGTGTTCCACA
GGGTAGCCAGCAGCATCCTGCGATGCAGATCCGGAACATAATGGTGCAGGGCGCTGACTTCCGCGTTT
CCAGACTTTACGAAACACGGAAACCGAAGACCATTCATGTTGTTGCTCAGGTCGCAGACGTTTTGCAG
CAGCAGTCGCTTCACGTTCGCTCGCGTATCGGTGATTCATTCTGCTAACCAGTAAGGCAACCCCGCCA
GCCTAGCCGGGTCCTCAACGACAGGAGCACGATCATGCGCACCCGTGGGGCCGCCATGCCGGCGATAA
TGGCCTGCTTCTCGCCGAAACGTTTGGTGGCGGGACCAGTGACGAAGGCTTGAGCGAGGGCGTGCAAG
ATTCCGAATACCGCAAGCGACAGGCCGATCATCGTCGCGCTCCAGCGAAAGCGGTCCTCGCCGAAAAT
GACCCAGAGCGCTGCCGGCACCTGTCCTACGAGTTGCATGATAAAGAAGACAGTCATAAGTGCGGCGA
CGATAGTCATGCCCCGCGCCCACCGGAAGGAGCTGACTGGGTTGAAGGCTCTCAAGGGCATCGGTCGA
GATCCCGGTGCCTAATGAGTGAGCTAACTTACATTAATTGCGTTGCGCTCACTGCCCGCTTTCCAGTC
GGGAAACCTGTCGTGCCAGCTGCATTAATGAATCGGCCAACGCGCGGGGAGAGGCGGTTTGCGTATTG
GGCGCCAGGGTGGTTTTTCTTTTCACCAGTGAGACGGGCAACAGCTGATTGCCCTTCACCGCCTGGCC
CTGAGAGAGTTGCAGCAAGCGGTCCACGCTGGTTTGCCCCAGCAGGCGAAAATCCTGTTTGATGGTGG
TTAACGGCGGGATATAACATGAGCTGTCTTCGGTATCGTCGTATCCCACTACCGAGATGTCCGCACCA
ACGCGCAGCCCGGACTCGGTAATGGCGCGCATTGCGCCCAGCGCCATCTGATCGTTGGCAACCAGCAT
CGCAGTGGGAACGATGCCCTCATTCAGCATTTGCATGGTTTGTTGAAAACCGGACATGGCACTCCAGT
CGCCTTCCCGTTCCGCTATCGGCTGAATTTGATTGCGAGTGAGATATTTATGCCAGCCAGCCAGACGC
AGACGCGCCGAGACAGAACTTAATGGGCCCGCTAACAGCGCGATTTGCTGGTGACCCAATGCGACCAG
ATGCTCCACGCCCAGTCGCGTACCGTCTTCATGGGAGAAAATAATACTGTTGATGGGTGTCTGGTCAG
AGACATCAAGAAATAACGCCGGAACATTAGTGCAGGCAGCTTCCACAGCAATGGCATCCTGGTCATCC
AGCGGATAGTTAATGATCAGCCCACTGACGCGTTGCGCGAGAAGATTGTGCACCGCCGCTTTACAGGC
TTCGACGCCGCTTCGTTCTACCATCGACACCACCACGCTGGCACCCAGTTGATCGGCGCGAGATTTAA
TCGCCGCGACAATTTGCGACGGCGCGTGCAGGGCCAGACTGGAGGTGGCAACGCCAATCAGCAACGAC
TGTTTGCCCGCCAGTTGTTGTGCCACGCGGTTGGGAATGTAATTCAGCTCCGCCATCGCCGCTTCCAC
TTTTTCCCGCGTTTTCGCAGAAACGTGGCTGGCCTGGTTCACCACGCGGGAAACGGTCTGATAAGAGA
CACCGGCATACTCTGCGACATCGTATAACGTTACTGGTTTCACATTCACCACCCTGAATTGACTCTCT
TCCGGGCGCTATCATGCCATACCGCGAAAGGTTTTGCGCCATTCGATGGTGTCCGGGATCTCGACGCT
CTCCCTTATGCGACTCCTGCATTAGGAAGCAGCCCAGTAGTAGGTTGAGGCCGTTGAGCACCGCCGCC
GCAAGGAATGGTGCATGCAAGGAGATGGCGCCCAACAGTCCCCCGGCCACGGGGCCTGCCACCATACC
CACGCCGAAACAAGCGCTCATGAGCCCGAAGTGGCGAGCCCGATCTTCCCCATCGGTGATGTCGGCGA
TATAGGCGCCAGCAACCGCACCTGTGGCGCCGGTGATGCCGGCCACGATGCGTCCGGCGTAGAGGATC
GAGATCGATCTCGATCCCGCGAAATTAATACGACTCACTATAGGGGAATTGTGAGCGGATAACAATTC
CCCTCTAGAAATAATTTTGTTTAACTTTAAGAAGGAGATATACATATGCACCATCATCATCATCATTC
TTCTGGTCTGGTGCCACGCGGTTCTGGTATGAAAGAAACCGCTGCTGCTAAATTCGAACGCCAGCACA
TGGACAGCCCAGATCTGGGTACCGACGACGACGACAAGGCCATGGCTGATATCGGATCCATGGCGACC
GAGGAGAATGTCAAGAGGCGAACACACAACGTCTTGGAGCGCCAGAGGAGGAACGAGCTAAAACGGAG
CTTTTTTGCCCTGCGTGACCAGATCCCGGAGTTGGAAAACAATGAAAAGGCCCCCAAGGTAGTTATCC
TTAAAAAAGCCACAGCATACATCCTGTCCGTCCAAGCAGAGACGCAAAAGCTCATTTCTGAAATCGAC
TTGTTGCGGAAACAAAACGAACAGTTGAAACACAAACTTGAACAGCTACGGAACTCTTGTGCGTAAGG
ACTCGAGCACCACCACCACCACCACTGAGATCCGGCTGCTAACAAAGCCCGAAAGGAAGCTGAGTTGG
CTGCTGCCACCGCTGAGCAATAACTAGCATAACCCCTTGGGGCCTCTAAACGGGTCTTGAGGGGTTTT
TTGCTGAAAGGAGGAACTATATCCGGATTGGCGAATGGGACGCGCCCTGTAGCGGCGCATTAAGCGCG
GCGGGTGTGGTGGTTACGCGCAGCGTGACCGCTACACTTGCCAGCGCCCTAGCGCCCGCTCCTTTCGC
TTTCTTCCCTTCCTTTCTCGCCACGTTCGCCGGCTTTCCCCGTCAAGCTCTAAATCGGGGGCTCCCTT

TAGGGTTCCGATTTAGTGCTTTACGGCACCTCGACCCCAAAAAACTTGATTAGGGTGATGGTTCACGT
AGTGGGCCATCGCCCTGATAGACGGTTTTTCGCCCTTTGACGTTGGAGTCCACGTTCTTTAATAGTGG
ACTCTTGTTCCAAACTGGAACAACACTCAACCCTATCTCGGTCTATTCTTTTGATTTATAAGGGATTT
TGCCGATTTCGGCCTATTGGTTAAAAAATGAGCTGATTTAACAAAAATTTAACGCGAATTTTAACAAA
ATATTAACGTTTACAATTTCAGGTGGCACTTTTCGGGGAAATGTGCGCGGAACCCCTATTTGTTTATT
TTTCTAAATACATTCAAATATGTATCCGCTCATGAATTAATTCTTAGAAAAACTCATCGAGCATCAAA
TGAAACTGCAATTTATTCATATCAGGATTATCAATACCATATTTTTGAAAAAGCCGTTTCTGTAATGA
AGGAGAAAACTCACCGAGGCAGTTCCATAGGATGGCAAGATCCTGGTATCGGTCTGCGATTCCGACTC
GTCCAACATCAATACAACCTATTAATTTCCCCTCGTCAAAAATAAGGTTATCAAGTGAGAAATCACCA
TGAGTGACGACTGAATCCGGTGAGAATGGCAAAAGTTTATGCATTTCTTTCCAGACTTGTTCAACAGG
CCAGCCATTACGCTCGTCATCAAAATCACTCGCATCAACCAAACCGTTATTCATTCGTGATTGCGCCT
GAGCGAGACGAAATACGCGATCGCTGTTAAAAGGACAATTACAAACAGGAATCGAATGCAACCGGCGC
AGGAACACTGCCAGCGCATCAACAATATTTTCACCTGAATCAGGATATTCTTCTAATACCTGGAATGC
TGTTTTCCCGGGGATCGCAGTGGTGAGTAACCATGCATCATCAGGAGTACGGATAAAATGCTTGATGG
TCGGAAGAGGCATAAATTCCGTCAGCCAGTTTAGTCTGACCATCTCATCTGTAACATCATTGGCAACG
CTACCTTTGCCATGTTTCAGAAACAACTCTGGCGCATCGGGCTTCCCATACAATCGATAGATTGTCGC
ACCTGATTGCCCGACATTATCGCGAGCCCATTTATACCCATATAAATCAGCATCCATGTTGGAATTTA
ATCGCGGCCTAGAGCAAGACGTTTCCCGTTGAATATGGCTCATAACACCCCTTGTATTACTGTTTATG
TAAGCAGACAGTTTTATTGTTCATGACCAAAATCCCTTAACGTGAGTTTTCGTTCCACTGAGCGTCAG
ACCCCGTAGAAAAGATCAAAGGATCTTCTTGAGATCCTTTTTTTCTGCGCGTAATCTGCTGCTTGCAA
ACAAAAAAACCACCGCTACCAGCGGTGGTTTGTTTGCCGGATCAAGAGCTACCAACTCTTTTTCCGAA
GGTAACTGGCTTCAGCAGAGCGCAGATACCAAATACTGTCCTTCTAGTGTAGCCGTAGTTAGGCCACC
ACTTCAAGAACTCTGTAGCACCGCCTACATACCTCGCTCTGCTAATCCTGTTACCAGTGGCTGCTGCC
AGTGGCGATAAGTCGTGTCTTACCGGGTTGGACTCAAGACGATAGTTACCGGATAAGGCGCAGCGGTC
GGGCTGAACGGGGGGTTCGTGCACACAGCCCAGCTTGGAGCGAACGACCTACACCGAACTGAGATACC
TACAGCGTGAGCTATGAGAAAGCGCCACGCTTCCCGAAGGGAGAAAGGCGGACAGGTATCCGGTAAGC
GGCAGGGTCGGAACAGGAGAGCGCACGAGGGAGCTTCCAGGGGGAAACGCCTGGTATCTTTATAGTCC
TGTCGGGTTTCGCCACCTCTGACTTGAGCGTCGATTTTTGTGATGCTCGTCAGGGGGGCGGAGCCTAT
GGAAAAACGCCAGCAACGCGGCCTTTTTACGGTTCCTGGCCTTTTGCTGGCCTTTTGCTCACATGTTC
TTTCCTGCGTTATCCCCTGATTCTGTGGATAACCGTATTACCGCCTTTGAGTGAGCTGATACCGCTCG
CCGCAGCCGAACGACCGAGCGCAGCGAGTCAGTGAGCGAGGAAGCGGAAGAGCGCCTGATGCGGTATT
TTCTCCTTACGCATCTGTGCGGTATTTCACACCGCATATATGGTGCACTCTCAGTACAATCTGCTCTG
ATGCCGCATAGTTAAGCCAGTATACACTCCGCTATCGCTACGTGACTGGGTCATGGCTGCGCCCCGAC
ACCCGCCAACACCCGCTGACGCGCCCTGACGGGCTTGTCTGCTCCCGGCATCCGCTTACAGACAAGCT
GTGACCGTCTCCGGGAGCTGCATGTG

**Table S1.** Overview of all synthesized proteins with their respective calculated and observed masses and rentention times.

| Miniprotein | Sequence | Yield (%) | Molecular weight calculated | Molecular weight observed | LCMS method | Retention time (min) | Observed ions |
|---|---|---|---|---|---|---|---|
| MiniMYC 1 | $(AcATEENVKRRTHNVLERQRRNELKRSFFG)_2K$ | 16% | 7137.1 | 3736.5 | B | 2.023 | 1190.3, 1020.4, 893.1, 793.9, 714.6, 649.7, 595.7, 549.9, 510.7, 476.7 |
| MonoMYC 2 | AcATEENVKRRTHNVLERQRRNELKRSCFA | 16% | 3483.0 | 3482.1 | B | 1.895 | 1161.5, 871.7, 697.5, 581.4, 498.5, 436.2 |
| MiniMYC 3 | $(AcATEENVKRRTHNVLERQRRNELKRSSCFA)_2$-Linker | 31% | 7260 | 7259.4 | B | 2.125 | 1210.9, 1038.1, 908.3, 807.5, 726.9, 660.9, 605.9, 559.4, 519.5 |
| MonoMYC 4 | AcATEENVKRRTHNVLERQRRNELKRSFFALRDQIPELENNEKAPKVVILKKATAYILSCA | 0.5% | 6978.1 | 6977.5 | B | 2.641 | 1396.6, 1163.8, 997.8, 873.1, 776.2, 698.7, 635.3, |
| DuoMYC 5 | $(AcATEENVKRRTHNVLERQRRNELKRSFFALRDQIPELENNEKAPKVVILKKATAYILSCA)_2$-linker | 23% | 14134 | 14133 | B | 2.897 | 1414.28, 1285.8, 1178.7, 1088.1, 1010.6, 943.2, 884.4, 832.4, 786.3, 744.9, 707.7, 674.0, 643.4 |
| MonoMYC 6 | AcVKRRTHNVLERQRRNELKRSFFALRDQIPELENNEKAPKVVILKKATAYILSVA$_\beta$C | 6% | 6532.7 | 6531.6 | B | 2.586 | 1089.5, 934.0, 817.4, 726.7, 649.0, 594.4 |
| DuoMYC 7 | $(AcVKRRTHNVLERQRRNELKRSFFALRDQIPELENNEKAPKVVILKKATAYILSA_\beta C)_2$-linker | 12% | 13359 | 13358 | B | 2.888 | 1114.0, 1028.5, 955.1, 891.1, 835.9, 786.8, 743.1, 704.1, 668.9, 637.1, 608.2, 581.7, 557.6 |
